# Chromosome-level genome assembly of the European Green woodpecker *Picus viridis*

**DOI:** 10.1101/2023.12.18.572251

**Authors:** Thomas Forest, Guillaume Achaz, Martial Marbouty, Amaury Bignaud, Agnès Thierry, Romain Koszul, Marine Milhes, Joanna Lledo, Jean-Marc Pons, Jérôme Fuchs

## Abstract

The European Green Woodpecker, *Picus viridis*, is a widely distributed species found in the Western Palearctic region. Here we assembled a highly contiguous genome assembly for this species using a combination of short and long reads sequencing and scaffolded with chromatin conformation capture (Hi-C). The final genome assembly was 1.28 Gb and features a scaffold N50 of 37Mb and a scaffold L50 of 39.165 Mb. The assembly incorporates 89.4% of the genes identified in birds in OrthoDB. Gene and repetitive content annotation on the assembly detected 15,805 genes and a ∼30.1% occurrence of repetitive elements, respectively. Analysis of synteny demonstrates the fragmented nature of the *Picus viridis* genome when compared to the chicken (*Gallus gallus*). The assembly and annotations produced in this study will certainly help for further research into the genomics of *P. viridis* and the comparative evolution of woodpeckers.

## Introduction

The European Green Woodpecker (*Picus viridis* Linnaeus, 1758) is a common Western Palearctic species that could be found in all sort of wooded environment where it often forages on ants on the ground. This species was previously considered to be conspecific with the Iberian Green Woodpecker (*Picus sharpei*) and Levaillant’s Woodpecker (*Picus vaillantii*) but recent molecular studies suggest that these taxa may consist of distinct biological species (Perktas et al. 2011, Pons et al. 2011, 2019). European Green and Iberian Green Woodpeckers form a secondary contact zone in southern France where individuals from the two species sometimes hybridize in the departments of Pyrénées-Orientales, Aude, Hérault and Gard, with very limited introgression outside of the contact zone (Pons et al. 2019). The European Green Woodpecker is currently considered as a common species (Least Concern) with a global increasing population size of 1.2 to 2.3 million individuals (BirdLife International, 2023), among which 300,000 – 600,000 are thought to occur in France (Deceuninck et al. 2015). The French population size is also considered to be increasing (+50% over the 1980-2012 time period, +2% over the 2001-2015 time period) (Deceuninck et al. 2015) and currently represents a stronghold of this species(25% of the total population).

Here we present a chromosome level assembly of the European Green Woodpecker genome, as well as genome resequencing data for another eleven individuals sampled over the 1970-2020 time period. Our assembly is based on a combination of Illumina short reads, Nanopore PromethION long reads and chromatin conformation capture (Hi-C). The vast majority of the *in silico* annotated genes are located on 47 chromosomal-level sequences, in accordance with the described karyotype of the species (2n=94, Hammar 1970).

## Material & Methods

### Sampling scheme

We sampled 13 *Picus viridis viridi*s individuals, five *historical* individuals collected between 1970 and 1977, and eight *contemporary* individuals collected between 2016 and 2021 (Table 1). Toe pads sampling for the historical specimens was performed using gloves and sterile scalpel blades, that were changed between each individual. Tissues for contemporary specimens consisted of muscle or heart either preserved in ethanol or flash frozen. For the two specimens used to assemble the reference genome, the tissues consisted of heart, muscle and liver (Nanopore long reads and Illumina short reads, MNHN ZO-2020-125) or liver (Hi-C, MNHN ZO-2021-2133) sampled 1-2 hours after the individual’s death and either immediately extracted (heart) or flash frozen at −80°C (liver). The two individuals originated from localities distant of 25 kms from each other and are both around 800 kms away from the secondary contact zone with its sister species *Picus sharpei* (Pons et al. 2019).

**Table 1.** Information of the sequenced individuals. All individuals were sampled in France.

| Specimen voucher | Date | Locality | Latitude | Longitude | Sex | Use |
| --- | --- | --- | --- | --- | --- | --- |
| **Historical** |  |  |  |  |  |  |
| ZO-1971-1090 | 16 Nov. 1970 | Gif-sur-Yvette, Essonne, Ile-de-France, France | 48.7 | 2.13 | M | Resequencing |
| ZO-1971-1091 | 28 Jul. 1971 | Baugé, Maine-et-Loire, Pays-de-la-Loire | 47.55 | -0.11 | M | Resequencing |
| ZO-1991-185 | 1972 | Chamonix-Mont-Blanc, Haute-Savoie, Auvergne-Rhône-Alpes | 45.93 | 6.93 | M | Resequencing |
| ZO-1991-186 | 1978 | Saclay, Essonne, Ile-de-France | 48.74 | 2.17 | F | Resequencing |
| ZO-1993-190 | 10 Apr. 1977 | Massérac, Loire-Atlantique, Pays-de-la-Loire | 47.67 | -1.91 | M | Resequencing |
| **Contemporary** |  |  |  |  |  |  |
| ZO-2017-195 | 7 Jun. 2016 | Vignely, Seine-et-Marne, Ile-de-France | 48.927248 | 2.805190 | F | Resequencing |
| ZO-2017-257 | 30 May 2016 | Évry-Grégy-sur-Yerre, Seine-et-Marne, Ile-de-France | 48.668917 | 2.651980 | F | Resequencing |
| ZO-2017-316 | 27 Jun. 2016 | Hagenthal-Le-Bas, Haut-Rhin, Grand-Est | 47.530303 | 7.482225 | M | Resequencing |
| ZO-2017-334 | 10 Jun. 2015 | Cavaillon, Vaucluse, Provence-Alpes-Côte-d'Azur | 43.836603 | 5.040671 | F | Resequencing |
| ZO-2018-033 | 15 Jun. 2016 | Niherne, Indre, Centre-Val-de-Loire | 46.772839 | 1.526087 | M | Resequencing |
| ZO-2018-044 | 18 Jul. 2016 | Blavozy, Haute-Loire, Auvergne-Rhône-Alpes | 45.062158 | 3.972074 | M | Resequencing |
| ZO-2020-125 | 1 Sep. 2020 | Courquetaine, Seine-et-Marne, Ile-de-France | 48.680485 | 2.749837 | M | Reference Genome<br>(Short/Long reads) |
| ZO-2021-2133<br>(fluid) | 12 Jul. 2021 | Serris, Seine-et-Marne, Ile-de-France | 48.857335 | 2.807369 | M | Reference Genome (Hi-C) |

### Extraction protocol

DNA was extracted using CTAB (resequencing, Winnepenninckx et al. 1993) or Phenol-Chloroform (reference genome) protocols. Historical specimens were extracted before any modern samples were processed. DNA quality and purity was assessed using a Qubit Quantification (Qubit dsDNA BR Assay Kit, Life Technologies), NanoDrop spectrophotometer (NanoDrop Technologies, Inc., Wilmington, DE) and Fragment Analyzer (Agilent) to assess DNA quality and quantity.

### Reference Genome sequencing

DNA-seq libraries have been prepared according to Illumina’s protocols using the Illumina TruSeq Nano DNA HT Library Prep Kit. Briefly, DNA was fragmented by sonication, size selection (average insert size : 400 bp) was performed using SPB beads (kit beads) and adaptators were ligated to be sequenced. Library quality was assessed using a Advanced Analytical Fragment Analyzer (Advanced Analytical Technologies, Inc., Iowa, USA) and libraries were quantified by qPCR using the Kapa Library Quantification Kit. DNA-sequencing experiments have been performed on a NovaSeq SP lane (Illumina, California, USA) using a paired-end read length of 2×150 pb with the Illumina NovaSeq Reagent Kits.

Long-reads library preparation was performed according to the manufacturer’s instructions ‘1D gDNA selecting for long reads (SQK-LSK109)’. At each step, DNA was quantified using the Qubit dsDNA HS Assay Kit (Life Technologies). DNA purity was tested using the nanodrop (Thermofisher) and size distribution and degradation assessed using the Fragment analyzer (Agilent) High Sensitivity Large Fragment 50kb Kit. Purification steps were performed using AMPure XP beads (Beckman Coulter). For one simple library, 10 µg of DNA was purified then sheared at 25kb using the megaruptor system (diagenode). A one step DNA damage repair + END-repair + dA tail of double stranded DNA fragments was performed on 2 µg of sample before ligating adapters to the library. Library was loaded onto one R9.4.1 flowcell at 20fmol then reloaded once at 11fmol on GridION instrument within 72H. For one optimised library, 20 µg of DNA was purified then sheared at 20kb using the megaruptor system (diagenode). A size selection step at 5kb using Short Read Eliminator XS Kit (Circulomics) was performed on 15µg of sample. For 5µg of DNA an extra repair step with SMRTbell DAMAGE REPAIR KIT (PACBIO, 100-465-900) was performed before the one step DNA damage repair + END-repair + dA tail of double stranded DNA fragments and adapters ligation. Library was loaded onto one R9.4.1 flowcell at 20fmol then reloaded three times at 20fmol on a PromethION instrument within 72H. All short and long-reads libraries were prepared and sequenced at the GeT-PlaGe core facility, INRAe Toulouse.

### Hi-C

Liver frozen tissue was directly put into 50 ml of Formaldehyde (3%) and Phosphate Buffered saline (PBS) solution. Fixation was then incubated for one hour under gentle agitation. Glycine was added to a final concentration of 0.125 mM and the reaction was incubated for another 20 minutes. Tissue was recovered by centrifugation (5,000g for 20 min) and washed with PBS 1X before being re-centrifuged. The Supernatant was discarded and tissue were stored at −80°C until use. The Hi-C library was constructed from liver tissue starting from a mass of 20 mg. Tissue was first resuspended in 1.2 mL of TE 1X and disrupted using CK14 glass beads (Precellys, Bertin Technology) and a precellys apparatus (program : 5 x 30 sec ON – 30 sec OFF – 8700 rpm). The - lysate was recovered and then use as input for the ARIMA-HiC preparation kit (Arima Genomics). Libraries were then processed for sequencing as previously described (Moreau et al. 2018) and sequenced on a Novaseq apparatus.

### Re-Sequencing of historical and contemporary samples

Re-sequencing of the eleven individuals was performed at the ‘Institut du Cerveau’ (ICM, Pitié-Salpêtrière Hospital, Paris) using a target 250-300 bp insert size on a NovaSeq 6000 system sequencer.

### Genome Reference assembly and quality assessment

A first assembly was made using the Nanopore long reads using the Flye (v2.8.1-b1676) *de novo* assembler (Kolmogorov et al. 2019). The assembly was then polished using three iterations of PILON (Walker et al. 2014), then once with Racon (Vaser et al. 2017), finished by a last iteration of PILON. The resulting polished assembly has been used as a guide for the Hi-C scaffolding process using instaGRAAL (Baudry *et al*. 2020).

### Genome size completeness, estimates of genome size using k-mer (SGA preqc)

SGA preqc tool (https://raw.githubusercontent.com/jts/sga/master/src/SGA/preqc.cpp) was used to do a k-mer analysis in order to estimate genome size completeness. K-mers, short DNA sequences of fixed length, were analyzed for their abundance in the genomic data. By calculating the occurrence of k-mers, it aims to make predictions about the size and completeness of the genome. It gives an hint on the quality of the data and anticipates the difficulty of the assembly process.

### K-mer composition of the genome assembly

To analyze the quality and composition of our genome assembly, we employed KAT (K-mer Analysis Toolkit) (Mapleson et al. 2017). This toolkit is designed for reference-free quality control of WGS (whole genome shotgun) reads and de novo genome assemblies. It utilizes k-mer frequencies and GC composition to assess errors, bias, and contamination throughout the assembly process. By comparing the k-mers present in the input reads and the resulting assemblies, it offers valuable insights into the composition and quality of genome assemblies.

### Masking repeated elements

We used RepeatMasker (v4.1.2-p1) (Smit et al. 2013) on the genome assembly obtained from the instaGRAAL step. This program screens DNA sequences for interspersed repeats and low complexity DNA sequences. It annotates the identified repeats and hard-masks them by replacing them with Ns.

### Genome annotation

The gene structure annotation was performed using BRAKER (Hoff et al. 2016), an automated method that utilizes both genomic and RNA-Seq data to generate comprehensive gene structure annotations in novel genomes.Using BRAKER-2.1.6, GeneMark (Lomsadze et al. 2005; Lomsadze et al. 2014) is first executed on a set of proteins coming from the reference proteome of *Gallus gallus* from UniProt (UP000000539). GeneMark is trained using these protein-coding sequences and produces a set of sequences for the *ab initio* methods used afterwards, like AUGUSTUS (Stanke et al. 2006), that produces gene and features predictions from the given *Picus viridis* genome. The resulting gene set consists of genes strongly supported by extrinsic evidence. Some statistics on the produced annotation have been generated using AGAT(v1.2.0) (Dainat et. Al 2023).

### Genome synteny

The masked genome was globally aligned with the genome of the chicken, *Gallus gallus*, the model organism, using MUMmer (Kurtz et al. 2004). We based our methodology for analyzing synteny on the one used for the reference genome assembly of *Colaptes auratus* (Hruska and Manthey, 2021). Only regions with more than 70% identity were retained, with lengths greater than 500bp. Next, the *Picus viridis* chromosomes showing the greatest overlap with chicken chromosomes were associated, and renamed according to their corresponding chicken chromosome name. A synteny circos plot produced with OmicCircos (v1.36) (Hu et al. 2014) shows the overall chromosome rearrangement between the two species. Only chromosomes sharing at least 500 unique links were conserved in the figure. To reorder chromosomes, the median position of hits in chickens for each segment from the *P. viridis* chromosomes was used to sort them. The same approach was performed against the genome of the Northern Flicker, *Colaptes auratus*, retaining fragments aligned with MUMmer (Kurtz et al. 2004) that exceed 10kb in length and exhibit 80% identity due to the significant noise resulting from close proximity and an abundance of repeated sequences.

### Assessing assembly quality

In order to estimate the quality we used BUSCO v4.1.4 to determine the proportion of genes expected in birds, using the Aves_ODB10 dataset, i.e. the 8,338 single-copy orthologous genes cataloged for the Aves class by the OrthoDB (v10) database (Simão et al. 2015).

### Ultra Conserved Elements

We also assessed genome completeness by estimating the number of Ultra Conserved Elements (UCEs), a set of loci commonly used in phylogenetics. We retrieved the UCEs from 13 Piciformes genomes published on Gebank (cut-off date 12 October 2022; Table S2). We extracted the UCEs with *Phyluce* (Faircloth 2016) according to the online tutorial (https://phyluce.readthedocs.io/en/latest/tutorials/tutorial-3.html). The UCEs from the 14 individuals were then aligned using MAFFT (Katoh et al. 2002) and internally trimmed with Gblocks (Castresana 2000; Talavera and Castresana 2007). We did not allow missing loci for a species. We performed a concatenated maximum likelihood analysis with the UCE loci under the GTRGAMMA model in RAXML (Stamatakis 2014).

**Table 2.**
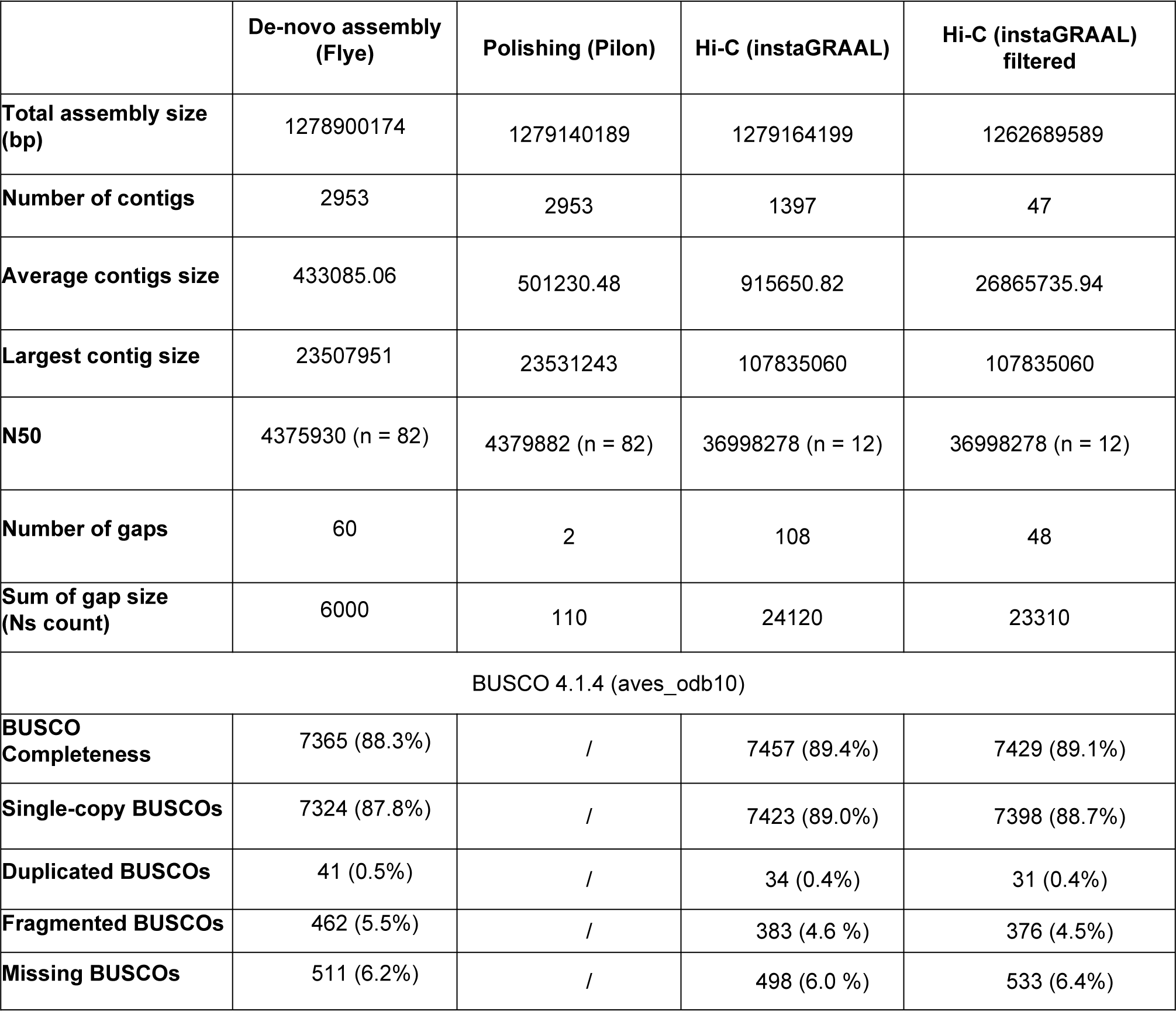
BUSCO analysis to assess the genome completeness. The analysis was performed three times, after significant filtering and assembly steps. Lines 1-7 of the table are given using Assembly Stats (v.1.0.1)

### Mitochondrial genome assembly

We used NOVOPlasty-2.7.2 (Dierckxsens et al. 2016) to assemble the mitochondrial genomes using the short reads. We specified a genome range of 16500-19000 bp, a K-mer size of k=39 and an insert size of 300 bp. As a seed, we used an ATP6 sequence from another *Picus viridis viridis* individual (MF766578, Shakya et al. 2017). When the genome was not circularized, we used the BWA algorithm, as implemented in Ugene (Okonechnikov et al. 2012) using the default option except the number of differences that we set to four.

We performed a mitochondrial genome analysis using all Piciformes sequences that are available on Genbank (cut-off date 15 July 2023). We restricted our analyses to 12 protein coding genes (we did not include ND6 as it was not present in several genbank sequences). We used the mitochondrial DNA genome of *Galbula dea* as an outgroup (MN356220) and included 36 other Piciformes (35 individuals available on Genbank and the newly produced sequence of *Picus viridis*) (Supplementary Table S1). We did not include *Yungipicus canicapillus* MK335534 because it is a chimeric sequence between *Yungipicus canicapillus* and *Dendrocopos darjellensis* (Fuchs et al, in preparation). Stop codons were excluded from the alignments as well as the extra nucleotide found in position 174 in ND3 in most Piciformes species. Substitution models were selected under the BIC criterion, as implemented in MEGA X (Kumar et al. 2018). Singe locus and concatenated partitioned Maximum Likelihood analyses were performed using RAXML on the RAXML Blackbox (Kozlov et al. 2019)(https://raxml-ng.vital-it.ch/). Clade robustness was estimated using one hundred bootstrap replicates.

### Population resequencing analysis and variant calling

From the newly assembled reference genome, we used snpArcher (Mirchandani et al. 2023) to call variants. This pipeline involves a mapping of all the reads on the reference genome and for those where the coverage and depth are sufficient, it marks each position that differs from the reference and merge all individuals into a single file in VCF format (Variant Call Format). We computed the number of segregating sites, S, and Watterson’s Θ, using PopGenome 2.7.5 (Pfeifer et al. 2014) and the Nei diversity index π (Nei and Li 1979) using VCFtools (Danecek et al. 2011) (Table 3). snpArcher automatically applies specific filters using the HaplotypeCaller from the Genome Analysis Toolkit (GATK) to customize our approach to the dataset’s characteristics. To heighten sensitivity and include variants in areas with inadequate read support, we set the --min-pruning filter to a permissive threshold of 1, meaning that variants with as little as one supporting read are considered for inclusion in the final variant call set. Additionally, the --min-dangling-branch-length filter is set to a value of 1, enabling even short ‘dangling branches’ in the assembly graph to be conserved and considering variants with minimal support. Nevertheless, it is essential to recognize that these filter settings boost sensitivity while potentially increasing the probability of false positive variant calls. These parameter selections were made carefully to achieve an equilibrium between sensitivity and specificity, while considering the characteristics of our data set. This has been carried out for subsequent filtering, after the construction of VCF, to enable fine-tuning of filters according to the observed depth distribution. Following these primary filters, further filtering based on coverage depth (DP) is applied to each variant, after the application of GATK caller filters. After implementing the initial filters, each variant undergoes depth-based filtering based on the depth of coverage (DP) once the GATK caller filters have been applied. Each variant is retained only if the depth exceeds 10X and falls below the 95th percentile of the observed depth distribution, in order to eliminate outliers.

**Table 3.** Summary statistics for the resequencing nuclear data, as estimated by PopGenome (Pfeifer et al. 2014) and VCFtools (Danecek et al. 2011).

|  | Total | Autosomes | Z chromosome |
| --- | --- | --- | --- |
| Number of segregating Sites (S) | 2407525 | 2103265 | 304260 |
| Normalized segregating sites by sequence length (s) | 0.001 | 0.001 | 0.002 |
| Nei diversity index ($\pi$) | 0.0009 | 0.0009 | 0.0008 |
| Watterson's estimator per site ($\Theta$) | 0.0005 | 0.0004 | 0.0007 |
| Historical individuals |  |  |  |
| Number of segregating Sites (S) | 1606510 | 1398591 | 207919 |
| Normalized segregating sites by sequence length (s) | 0.001 | 0.001 | 0.001 |
| Nei diversity index ($\pi$) | 0.0011 | 0.0011 | 0.0010 |
| Watterson's estimator per site ($\Theta$) | 0.0004 | 0.0004 | 0.0006 |
| Contemporary individuals |  |  |  |
| Number of segregating Sites (S) | 1606510 | 990064 | 145932 |
| Normalized segregating sites by sequence length (s) | 0.001 | 0.0008 | 0.001 |
| Nei diversity index ($\pi$) | 0.0008 | 0.0009 | 0.0007 |
| Watterson's estimator per site ($\Theta$) | 0.0004 | 0.0002 | 0.0004 |

### Site Frequency Spectrum

The Site Frequency Spectrum (SFS) represents the distribution of the allelic frequencies of the mutations throughout the genome (Fu 1995). It gives the number of mutations present at each frequency. The folded SFS of a sample of n diploid individuals is described as a vector *η* such that *η* = (*η*_1_, *η*_2_, …, *η*_2n−1_), where *η*_i_ is the number of mutations at frequency i/2n with i∈[1 : 2_n_ − 1]. Spectra are also normalized and transformed to *ϕ*_i_ = *η*_i_ × i(n − I)/Σ*η*_i_, except for i = n where *ϕ*_i_ = n/2Σ*η*_i_ (Achaz, 2009).

## Results

### Reference Genome sequencing

Heart provided less degraded DNA when compared to pectoral muscle or liver for long-read sequencing. We obtained 41.2 Gbp of long reads (Gridion: 3.5 Gbp, N50 close to 16 kb; Promethion: 37.7 Gbp, N50 close to 10 kb) as well as about 65 Gbp of short-read data. The SGA preqc genome size estimate for *P. viridis* was 1.26 Gb. From mapping information, coverage depth is around 73X. (Table S5)

**Figure 1.**
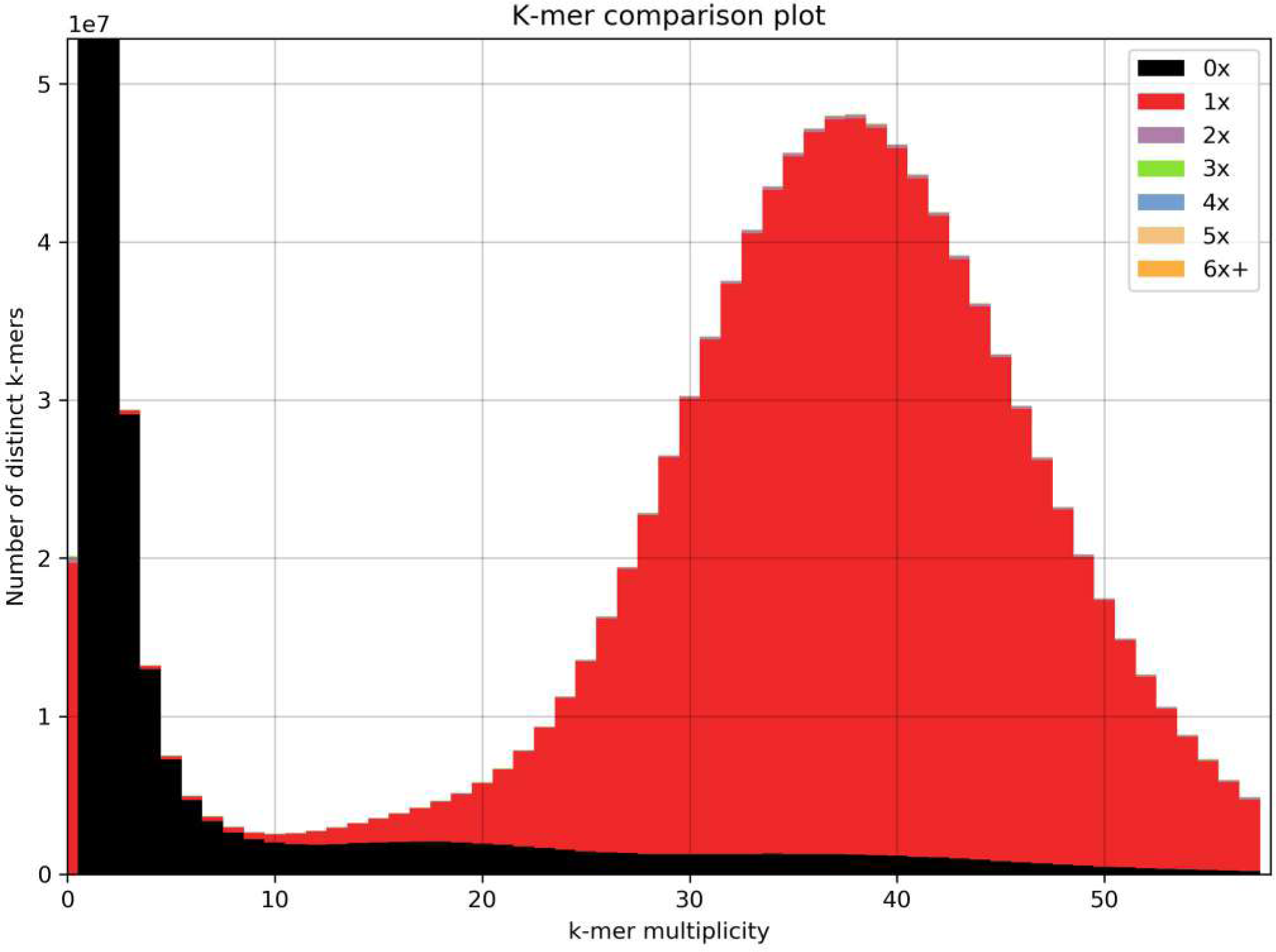
Distribution of k-mers using KAT. In the k-mer spectrum, reads that are absent from the assembly are displayed in black. The error distribution is quite low with most of errors under 10x.

### Assembly quality and genome completness

The majority of the assembly presents unique k-mer content, that is only present once. This pattern is typical of what we expect from a complete haploid assembly, generated from a diploid genome. Results from the BUSCO analyses indicated that 93.6% of the loci were included in the assembly (single-copy: 88.7%, duplicated: 0.4%, fragmented: 4.5%). Comparison with other Picidae genomes indicated our genome assembly ranks third out of five for genome completeness (Table S2) and third out of four for genomes for which long reads were generated.

The second highest number of retrieved UCEs in the Piciformes was found in our *P. viridis* assembly, confirming its high completeness (Supplementary Table S3). Alignment of the concatenated UCEs (2906 loci) was 631,268 bp long, among which 16,077 sites were informative. The result of the Maximum Likelihood analyses recovered the same topology (Figure 2) at the family level than Prum et al. (2015), with all nodes being supported by bootstrap values of 100.

**Figure 2.**
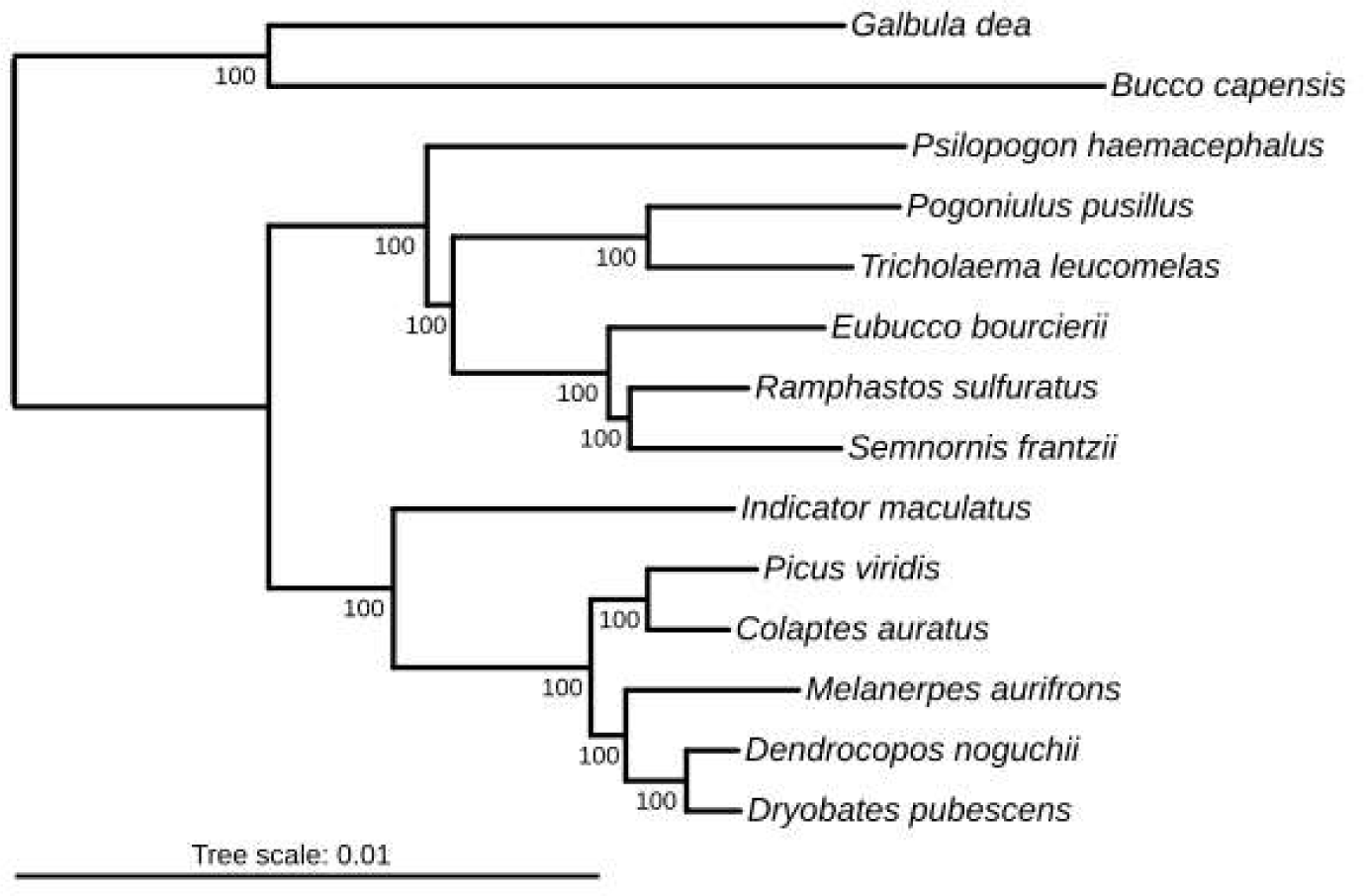
Phylogenetic tree showing relationships among Piciformes species resulting from maximum likelihood analysis of Ultra Conserved Elements loci. The tree was reconstructed from an alignment of 631,268 nucleotides (2906 loci). The bootstrap percentage (BP) is indicated on the nodes.

### Mitogenome assembly

Our final assembly of the mitochondrial genome was 16,912 bp long, with some uncertainty concerning the exact length due to the presence of a 64 bp repeat at the end of the first control region as well as one C monomeric region in 16S (Table S4). The genome is similar to many other avian genomes with 13 protein coding genes, 22 transfert RNA and 2 Ribosomal RNA. The control region is duplicated, presenting one functional control region and one degenerated control region.

The topology recovered from the partitioned concatenated analysis of twelve mitochondrial protein coding loci (Figure 3) was in strong agreement with current phylogenetic hypotheses for family-level and genus-level relationships in Piciformes (e.g. Prum et al. 2015, Shakya et al. 2017).

**Figure 3.**
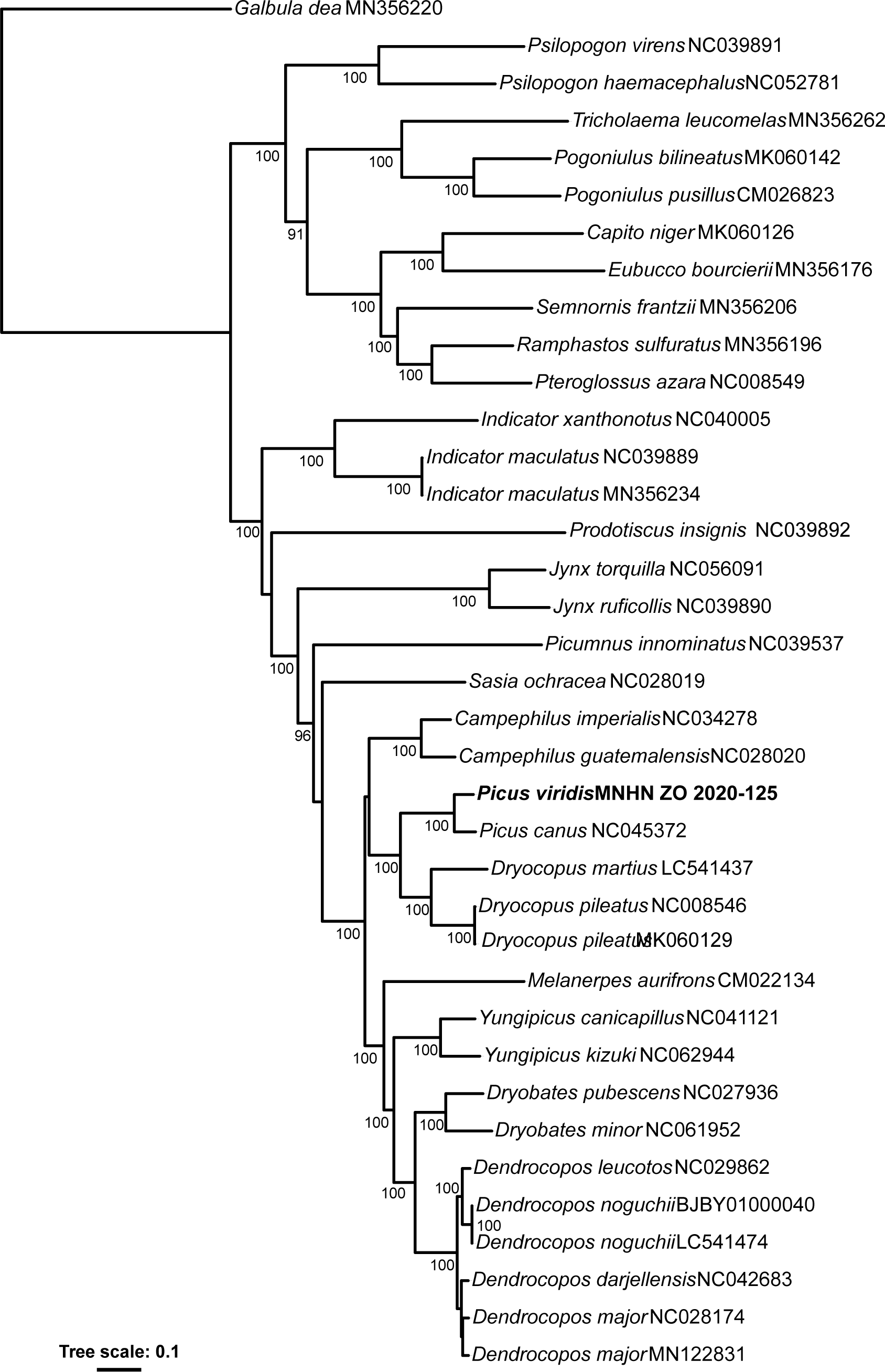
Maximum Likelihood phylogenetic tree showing relationships among Piciformes species. Twelve mitochondrial loci, 37 taxa and 10,857 characters were included in the analysis. The bootstrap percentage (BP) is indicated on the nodes.

### Genome annotation

A total of 15,848 protein-coding genes were identified, representing 81357234bp in total, along with essential transcription factors, non-coding RNAs, and repetitive elements, through automated genome annotation. 16212 mRNA, representing 82809862bp in total, were identified, including 364 isoforms. 77423 exons are numbered, at a rate of 4.8 exons per Coding DNA Sequence. (Table S9)

### Repetitive elements

Detailed analysis of repetitive elements in the *Picus viridis* genome revealed their abundance, comprising 30.1 % of the assembled genome (385325029 out of 1279164199 total bp). This estimate compares with the 28% recovered for *Colaptes auratus* (Hruska and Manthey 2021), 25.8% for *Melanerpes aurifrons* and 22% for *Dryobates pubescens* (Zhang et al. 2014). Details of the types of repeated elements found using Repeatmasker can be found in the supplementary material. (Table S8)

### Genome Synteny

Two circos plots were created to compare the genome assembly produced for *Picus viridis* against the chicken, *Gallus gallus*, (Figure 4) and the Northern Flicker, *Colaptes auratus* (Figure 5), respectively. *Gallus gallus* was used as a reference because it is commonly employed as a model organism for comparative purposes. Furthermore, their repeat element rate is reflective of that of most avian species (∼10%) (Hillier et al. 2004). The European Green Woodpecker genome exhibits higher fragmentation than that of the chicken and is more similar to the genome of *C. auratus* in terms of fragmentation, with an abundance of micro-chromosomes. This similarity is noteworthy and may accounts for the difficulties encountered in identifying expected genes in databases such as OrthoDB which logically reduces the BUSCO completeness score.

**Figure 4.**
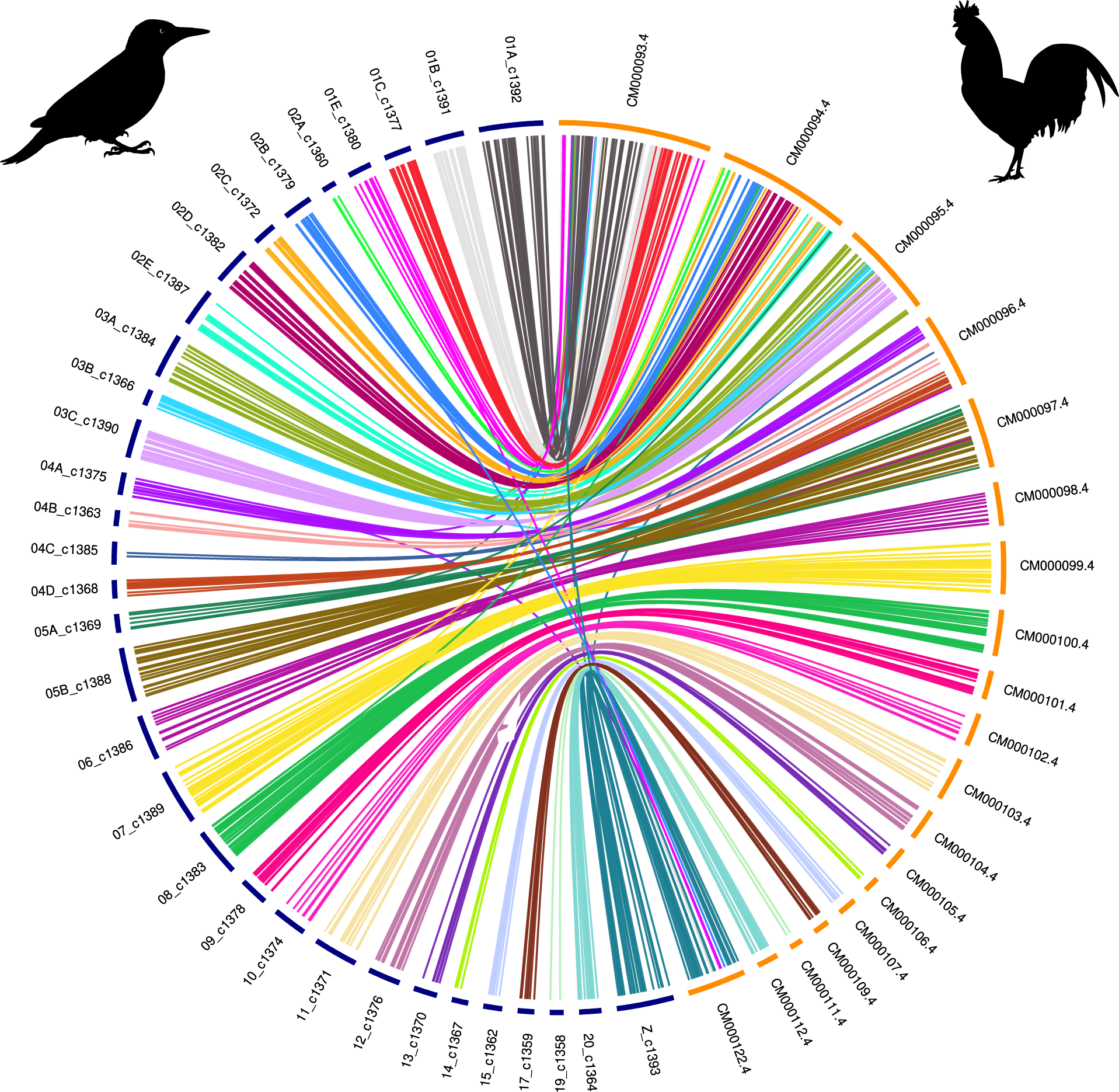
Circos plot. Woodpecker chromosomes are on the left, in blue; chicken chromosomes are on the right, in yellow. Note that the number of chromosomes differs between the Chicken *Gallus gallus* and the European Green Woodpecker *Picus viridis*.

**Figure 5.**
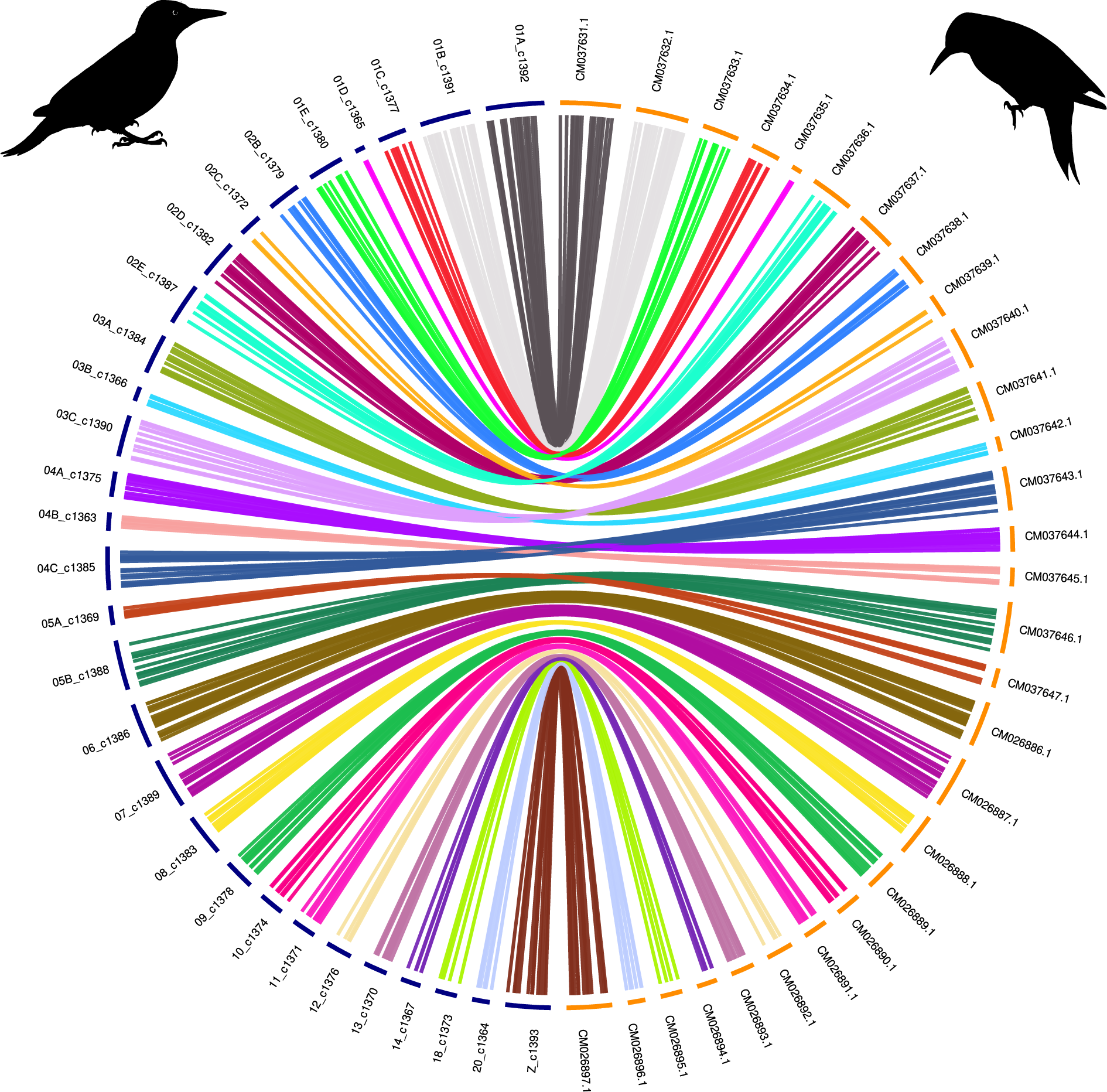
Circos plot. Woodpecker chromosomes are on the left, in blue; Northern flicker (*Colaptes auratus*) chromosomes are on the right, in yellow.

### Nuclear diversity

In the final VCF, variants with coverage depth within the [10;346] interval are retained, where 346X represents the value above which 90% of reads are observed in the depth distribution. Before applying depth-based filters, 8,815,726 variants were identified among the 12 individuals. The average depth of variants is 22.7X. We retain 7,226,988 variants if the reads fall within the [10; 346]X range. (346X is the 90th percentile of the depth distribution). Higher coverage values are likely caused by repeated sequences Genetic diversity values were higher in historical specimens than in contemporaneous. The Z chromosome was comparatively more diverse than autosomes (Table 3). On both the Principal Component Analysis (PCA) (Figure 6) and the hierarchical clustering dendrogram using a complete-linkage method (Figure 7), based on sequence dissimilarities, historical and contemporaneous individuals do appear to cluster into two groups. These two groups can be formed by cutting the dendrogram at a dissimilarity value of 1.0. On the PCA, historical individuals appear slightly more genetically diverse. One historical individual (ZO-1971-1090) is however nested among the contemporary individuals. Frequency spectra (Figure 8) suggest that the population does not have a constant panmictic demographic history. This pattern may be due to changes in population structure and/or significant changes in population size in the past that are compatible with an expanding population, characterized by an excess of low frequency variants.

**Figure 6.**
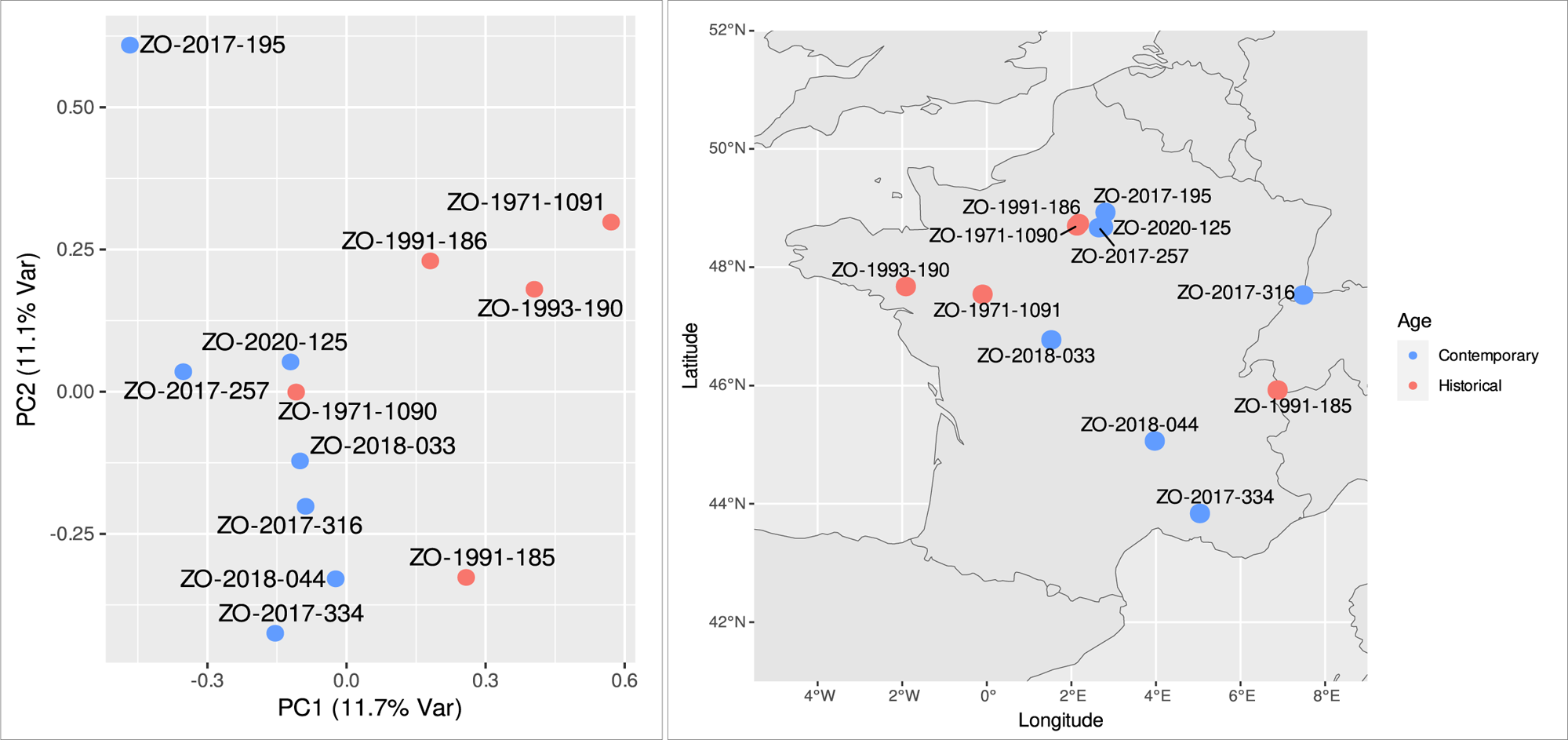
Principal Component Analysis based on genomic diversity among the historical and contemporary samples (left) and map of the sampled *Picus viridis* individiduals (right).

**Figure 7.**
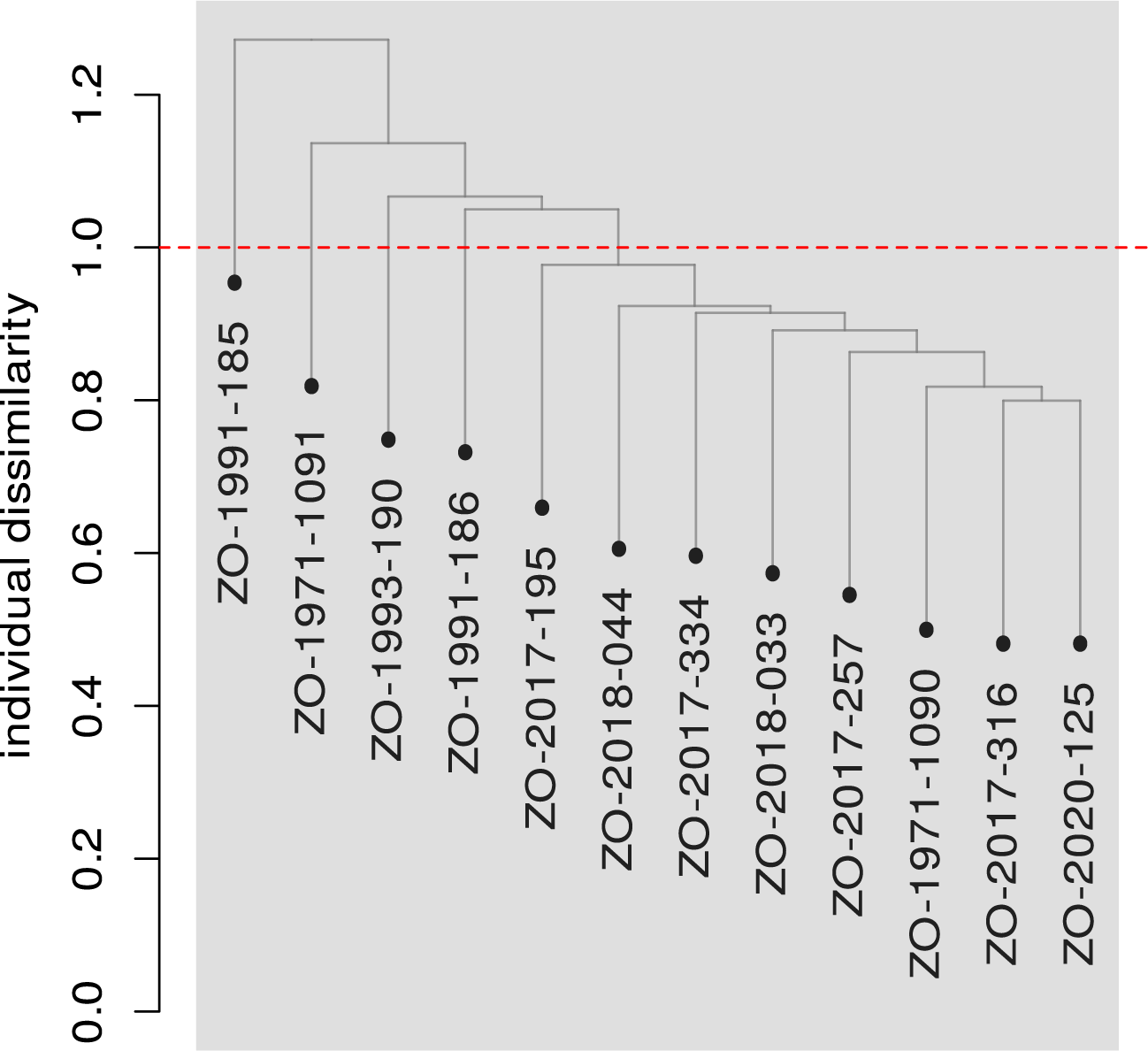
Dendrogram of hierarchical clustering based on pairwise nuclear sequence dissimilarity between samples.

**Figure 8.**
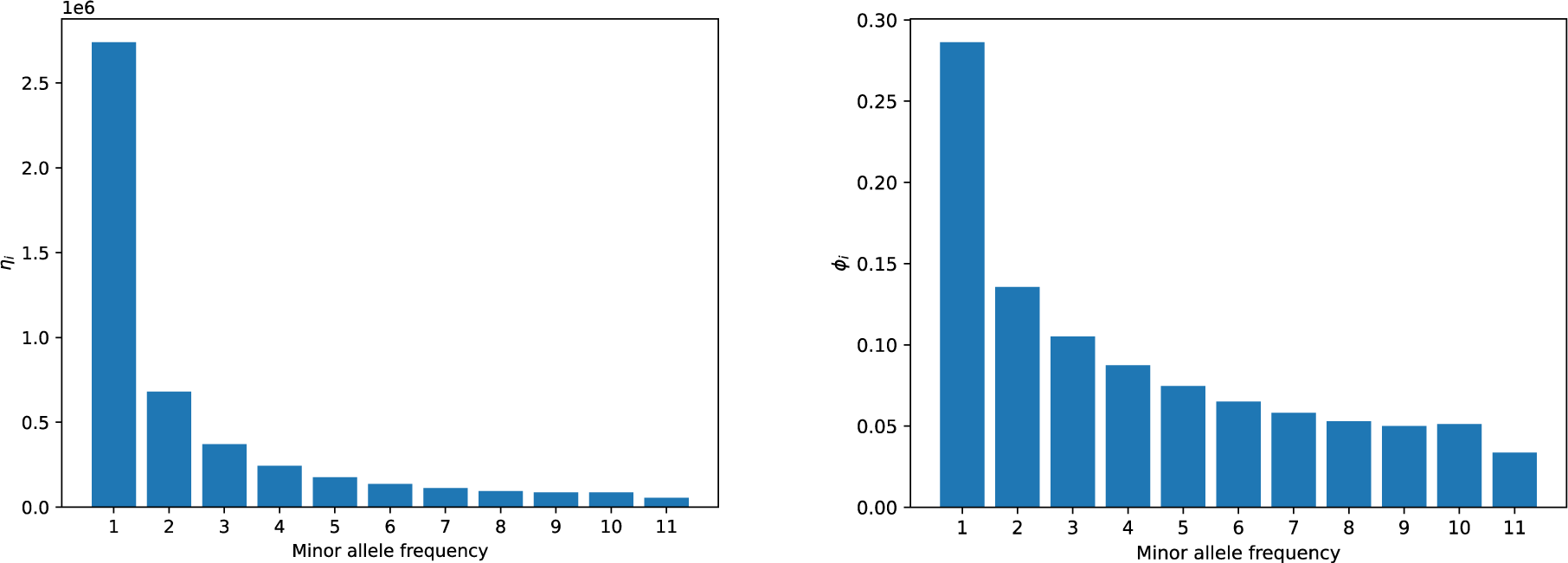
Site Frequency Spectrum. On left, the raw counts of each allele frequency, on the right, the normalized (rescaled [0,1]) and transformed, so that the expected spectrum for a constant population size should be flat at y=1/11.

### Mitogenome diversity

We obtained circularized assemblies for five out of the twelve samples; we observed length variation among the mitochondrial genomes assembled by Novoplasty, mostly due to the number of a 64 bp repeat at the end of the first control region (which varies between two and five). Two regions were difficult to handle by the algorithm using the shorts reads: the aforementioned 64 bp region located in Control Region 1 as well as one monomeric region in 16S that consists of up to 37 C. Similar problems were found with the BWA mapping algorithm, although the latter tended to provided short genomes (the 64 bp repeat being usually considered as a single repeat with an increase in estimated coverage).

Prior to the analyses, we excluded three regions (positions 966-977 in 12S, position 1538-1574 in 16S and 16023-16524 in Control Region) that either consist of monomeric C regions (position 966-977 and 1538-1574) or of the 64 bp region with uncertain number of repeats, resulting in a final alignment of 16769 bp.

Summary statistics (number of haplotypes H, haplotype diversity Hd, number of segregating sites S, nucleotide diversity π, Watterson’s theta Θ), as estimated by DNAsp 5 (Librado et al. 2009) are reported in Table 4. As well as the nuclear data, genetic diversity values are higher in historical specimens than in contemporaneous specimens. The minimum spanning network, as estimated by Pegas (Paradis 2010) is shown in Figure S1. The average p-distance among individuals is 0.11% (minimum: 0.005% between two contemporary individuals sampled 400 kms apart and 0.17% between a contemporary and historical individual sampled 400 kms apart as well).

**Table 4.** Summary statistics for the mitochondrial data, as estimated using DNAsp 5 (Librado et al. 2009)

|  | **All** | **Historical (n=5)** | **Contemporary (n=7)** |
| --- | --- | --- | --- |
| **H** | 12 | 5 | 7 |
| **Hd** | 1 | 1 | 1 |
| **S** | 87 | 44 | 48 |
| **$\pi$** | 0.00113 | 0.00115 | 0.00104 |
| **$\Theta$** | 0.00172 | 0.00126 | 0.00117 |

## Discussion

Using a combination of short and long reads as well as HiC data, we successfully assembled a reference genome for a species that had no existing reference genome. The final genome assembly of European Green Woodpecker is estimated to be about 1.28 Gbp. Exhibiting an N50 contig length of ∼37 Mbp, our assembly displays significant contig continuity, retaining almost all genetic information across the 47 contiguous scaffolds. The de novo assembly and annotation of the *Picus viridis* genome presented here represent an important resource for the ornithological community and complements our understanding of the genetic structure of the European Green Woodpecker’s. but it is important to acknowledge that challenges persist with any genome assembly (Peona et al. 2018; Weissensteiner and Suh 2019). Although continuity and completeness have significantly improved, some genomic regions with high GC content and repetitive elements may still present challenges for accurate assembly (Chen et al. 2013; Goldstein et al. 2019).

### Genome completeness

The BUSCO completeness score of 89.4% (Table 2) is comparatively low for recently assembled bird genomes, especially considering the methods used (Hi-Ci, long reads, and short reads) when compared to other recently published genomes (e.g. Baudrin et al. 2023). Among Piciformes, our assembly ranked third out of four species (*Melanerpes aurifrons, Dryobates pubescens, Picus viridis, Colaptes auratus*) for which long-reads were used. We note, however that the two assemblies with the lowest BUSCO scores (*Colaptes auratus*, *Picus viridis*) are part of the Picini subclade whereas the two higher scores belong to the Dendropicini (*Dryobates pubescens*) and Melanerpini (*Melanerpes aurifrons*) clades (Shakya et al. 2017). Genome fragmentation, especially the number of microchromosomes, could explain this pattern as they could be more difficult to assemble. Yet, karyotypic data indicate that *Dryobates pubescens* (2n=92, Shields et al. 1982), *Colaptes auratus* (2n=90, Shields et al. 1982) *Picus viridis* (2n=94; Hammar 1970) have similar number of chromosomes. The karyotype of *Melanerpes aurifrons*, although currently unknown is possibly much lower as one of its congener, *M. candidus*, has a chromosome number of 2n=64 (de Oliveira et al. 2017). Woodpeckers, unlike most other birds, are particularly rich in repetitive elements (Zhou et al. 2014; Manthey et al. 2018). The proportion of repetitive elements is a strong particularity, among birds, of the Piciformes. The proportion of the genome that consists of repetitive elements is the highest in the two species (*Picus viridis*: 30.1%, *Colaptes auratus*: 28%) that have the lower BUSCO scores. It is plausible that the high proportion of repetitive elements in the genomes could also contribute to the lower recovery of BUSCO loci in the assemblies.

### Spatial structuration in historical and contemporaneous data

On the PCA (Figure 6), there is a visible genomic structure following a north-south gradient on PC2, and a temporal differentiation based on PC1. Indeed the genomic information overall indicates a genetic proximity to birds from nearby regions. Yet, one individual, ZO-2017-195, from the Ile-de-France area, where several modern samples were sequenced, appears one outlier individual on the PCA, although it is located more internally in the dendrogramm (Figure 7). Temporal genetic structuring may exist, where current (expanding) populations are the result of expansion of only a subset of the 1970-1980 population, with potential replacement. Indeed, population expansion may not be homogenous and it is conceivable that not all subpopulations contributed equally to the strong populations expansion documented in the last 50 years. Population genomic studies from different time periods will be needed to test this hypothesis.

## Data Availability

The PicVir_MNHN_1.0 assembly can be accessed at NCBI (BioProject PRJNA1027323; Genome GCA_033816785.1). All of the related raw sequencing data, specifically Illumina, Nanopore, and Hi-C, can be obtained through NCBI SRA under the same BioProject. Scripts, associated files for this project can be found on GitHub (https://github.com/tforest/P_viridis_assembly). Supplementary files containing data from BUSCO, BRAKER, outputs of MUMmer alignments, genome annotation in GFF format, the VCF file and supplementary material are available on figshare: https://figshare.com/s/e7562104b84a0c6e6e89.

## Acknowledgments

We are very grateful to the multiple wildlife rehabilitation centers: ENVA/CHU-FS (Pierrick Poiré Cécile Le Barzic, Jean-François Courreau, Pascal Arné) and the Centres de soins LPO (Brenne, Tony Williams; Buoux, Alexandra de Kerviler, Aurélie Amiault; Clermont-Ferrand, Adrien Corsi; Rosenwiller, Emilie Dusausoy, Suzel Hurstel, Jade Oliva) for their collaboration on our salvage bird program. DNA extraction and quantification were performed at the Service de Systématique Moléculaire (UAR 2700 2AD), and we acknowledge the help of its staff during the laboratory work. This work was performed in collaboration with the GeT core facility, Toulouse, France. The Hi-C work was possible thanks to the team of Spatial Regulation of Genomes at the Pasteur Institute in Paris. Most of the bioinformatic analyses were carried out through the Genotoul cluster, except for the variant calling process through the snpArcher pipeline, which was executed on the PCIA cluster (Plateforme de Calcul Intensif et Algorithmique PCIA, Muséum national d’histoire naturelle, Centre national de la recherche scientifique, UAR 2700 2AD).

---

**Figure S1.**
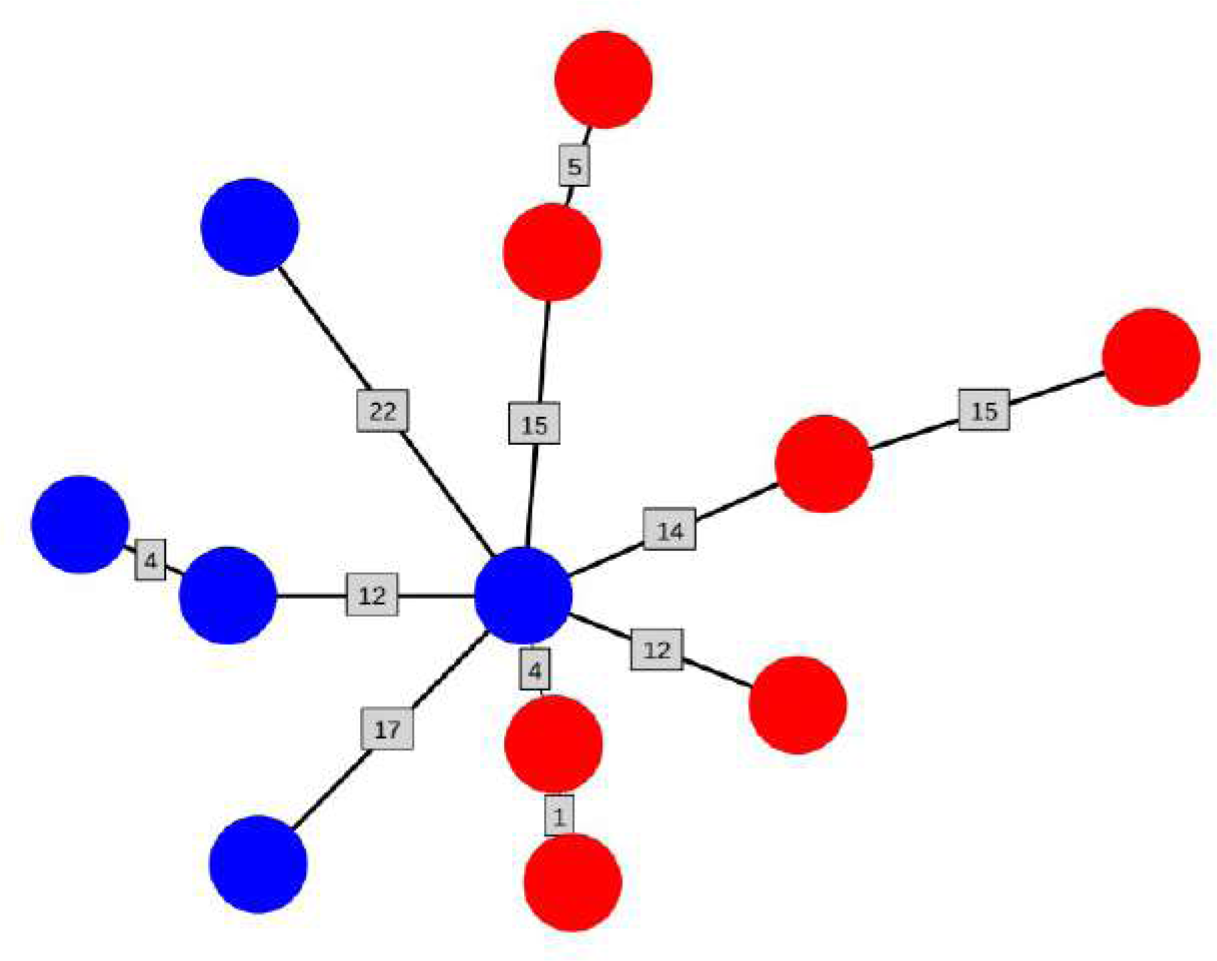
Mitochondrial minimum Spanning Network built using Pegas (Paradis 2010). Colors refer to historical (blue) and contemporary (red) samples. Number in squares represent the number of mutations.

**Table S1.** Sampling scheme for the mitogenome analysis.

| Family | Genera | Species | GenBank Accession Number | Reference |
| --- | --- | --- | --- | --- |
| Capitonidae | *Capito* | *niger* | MK060126 | Tamashiro et al. (2018) |
| Galbulidae | *Galbula* | *dea* | MN356220 | Feng et al. (2020) |
| Indicatoridae | *Indicator* | *maculatus* | NC039889 | Tamashiro et al. (2018) |
| Indicatoridae | *Indicator* | *maculatus* | MN356234 | Feng et al. (2020) |
| Indicatoridae | *Indicator* | *xanthonotus* | NC040005 | Duan et al. (2018) |
| Indicatoridae | *Prodotiscus* | *insignis* | NC039892 | Tamashiro et al. (2018) |
| Lybiidae | *Pogoniulus* | *bilineatus* | MK060142 | Tamashiro et al. (2018) |
| Lybiidae | *Pogoniulus* | *pusillus* | CM026823 | Kirschel et al. (unpublished) |
| Lybiidae | *Tricholaema* | *leucomelas* | MN356262 | Feng et al. (2020) |
| Megalaimidae | *Psilopogon* | *virens* | NC039891 | Tamashiro et al. (2018) |
| Megalaimidae | *Psilopogon* | *haemacephalus* | NC052781 | Feng et al. (2020) |
| Picidae | *Campephilus* | *guatemalensis* | NC028020 | Fuchs et al. (2015) |
| Picidae | *Campephilus* | *imperialis* | NC034278 | Anmarkrud and Lifjeld JT (2017) |
| Picidae | *Dendrocopos* | *darjellensis* | NC042683 | Bi et al. (2019) |
| Picidae | *Dendrocopos* | *leucotos* | NC029862 | Eo (2017) |
| Picidae | *Dendrocopos* | *major* | NC028174 | Park et al. (2019) |
| Picidae | *Dendrocopos* | *major* | MN122831 | Unpublished |
| Picidae | *Dryobates* | *pubescens* | NC027936 | Zhang et al. (2016) |
| Picidae | *Dryobates* | *minor* | NC061952 | Chen et al. (2022) |
| Picidae | *Dryocopus* | *martius* | LC541437 | Yamamoto et al. (unpublished) |
| Picidae | *Dryocopus* | *pileatus* | NC008546 | Gibb et al. (2007) |
| Picidae | *Dryocopus* | *pileatus* | MK060129 | Tamashiro et al. (2018) |
| Picidae | *Jynx* | *ruficollis* | NC039890 | Tamashiro et al. (2018) |
| Picidae | *Jynx* | *torquilla* | NC056091 | Du et al. (2020) |
| Picidae | *Melanerpes* | *aurifrons* | CM022134 | Wiley and Miller (2020) |
| Picidae | *Picumnus* | *innominatus* | NC039537 | Zhou et al. (2017) |
| Picidae | *Picus* | *canus* | NC045372 | Yao et al. (2019) |
| Picidae | *Picus* | *viridis* | To be added | This study |
| Picidae | *Dendrocopos* | *noguchii* | BJBY01000040.1 | Nakajima et al. (unpublished) |
| Picidae | *Dendrocopos* | *noguchii* | LC541474 | Yamamoto et al. (unpublished) |
| Picidae | *Sasia* | *ochracea* | NC028019 | Fuchs et al. (2015) |
| Picidae | *Yungipicus* | *canicapillus* | NC041121 | Lai et al. (2019) |
| Picidae | *Yungipicus* | *kizuki* | NC062944 | Kim (unpublished) |
| Ramphastidae | *Eubucco* | *bourcierii* | MN356176 | Feng et al. (2020) |
| Ramphastidae | *Pteroglossus* | *azara* | NC008549 | Gibb et al. (2007) |
| Ramphastidae | *Ramphastos* | *sulfuratus* | MN356196 | Feng et al. (2020) |
| Ramphastidae | *Semnornis* | *frantzii* | MN356206 | Feng et al. (2020) |

**Table S2.**
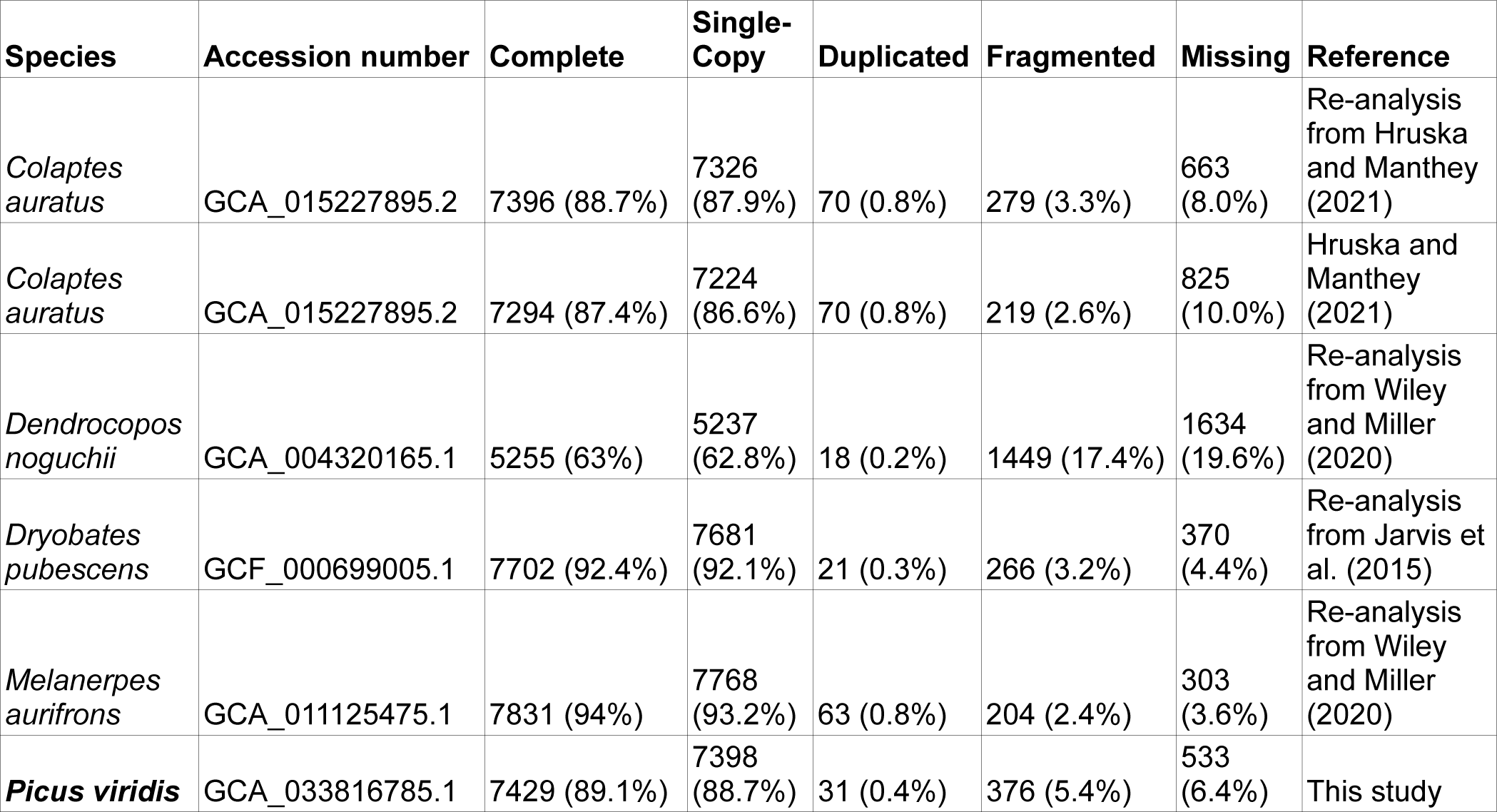
BUSCO scores among six Picidae species.

| Species | Accession number | Complete | Single-Copy | Duplicated | Fragmented | Missing | Reference |
| --- | --- | --- | --- | --- | --- | --- | --- |
| *Colaptes auratus* | GCA_015227895.2 | 7396 (88.7%) | 7326 (87.9%) | 70 (0.8%) | 279 (3.3%) | 663 (8.0%) | Re-analysis from Hruska and Manthey (2021) |
| *Colaptes auratus* | GCA_015227895.2 | 7294 (87.4%) | 7224 (86.6%) | 70 (0.8%) | 219 (2.6%) | 825 (10.0%) | Hruska and Manthey (2021) |
| *Dendrocopos noguchii* | GCA_004320165.1 | 5255 (63%) | 5237 (62.8%) | 18 (0.2%) | 1449 (17.4%) | 1634 (19.6%) | Re-analysis from Wiley and Miller (2020) |
| *Dryobates pubescens* | GCF_000699005.1 | 7702 (92.4%) | 7681 (92.1%) | 21 (0.3%) | 266 (3.2%) | 370 (4.4%) | Re-analysis from Jarvis et al. (2015) |
| *Melanerpes aurifrons* | GCA_011125475.1 | 7831 (94%) | 7768 (93.2%) | 63 (0.8%) | 204 (2.4%) | 303 (3.6%) | Re-analysis from Wiley and Miller (2020) |
| ***Picus viridis*** | GCA_033816785.1 | 7429 (89.1%) | 7398 (88.7%) | 31 (0.4%) | 376 (5.4%) | 533 (6.4%) | This study |

**Table S3.** Information on sequences used in the UCE analysis.

| **Species** | **Accession Number** | **Number of UCE loci retrieved - out of 5041** | **Reference** |
| --- | --- | --- | --- |
| *Galbula dea* | GCA_013399015.1 | 4811 | Feng et al. (2020) |
| ***Picus viridis*** | **GCA_033816785.1** | **4742** | **This study** |
| *Indicator maculatus* | GCA_013399975.1 | 4722 | Feng et al. (2020) |
| *Dendrocopos noguchii* | GCA_004320165.1 | 4707 | Unpublished |
| *Tricholaema leucomelas* | GCA_013400475.1 | 4690 | Feng et al. (2020) |
| *Pogoniulus pusillus* | GCA_015220805.1 | 4684 | Kirschel et al. (2021) |
| *Ramphastos sulfuratus* | GCA_013399055.1 | 4676 | Feng et al. (2020) |
| *Bucco capensis* | GCA_013396975.1 | 4672 | Feng et al. (2020) |
| *Dryobates pubescens* | GCA_000699005.1 | 4664 | Zhang et al. (2014) |
| *Melanerpes aurifrons* | GCA_011125475.1 | 4662 | Wiley and Miller (2020) |
| *Eubucco bourcierii* | GCA_013396675.1 | 4657 | Feng et al. (2020) |
| *Psilopogon haemacephalus* | GCA_013396835.1 | 4571 | Feng et al. (2020) |
| *Colaptes auratus* | GCA_015227895.2 | 4063 | Hruska et a. (2021) |
| *Semnornis frantzii* | GCA_013399775.1 | 4044 | Feng et al. (2020) |

**Table S4.**
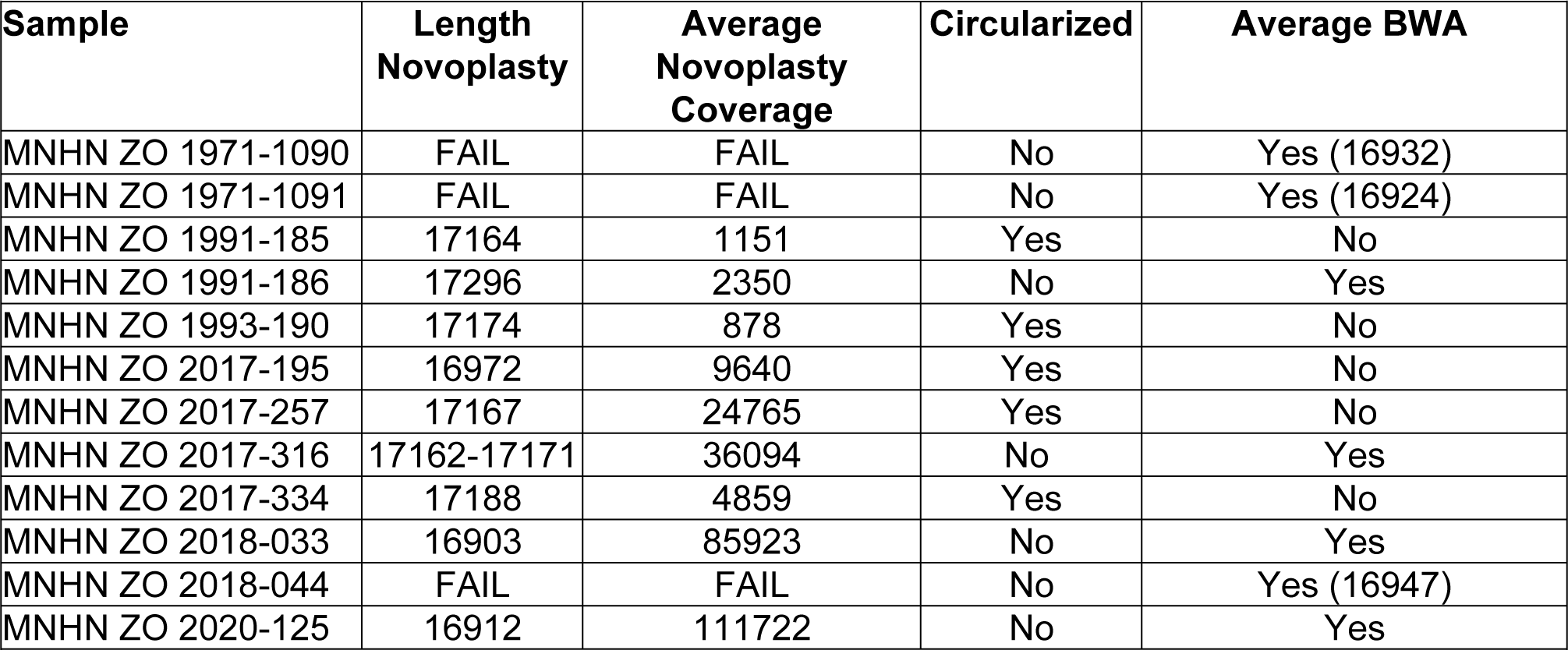
Mitochondrial genome as assembled by Novoplasty and BWA. The BWA mapping is not performed for circularized genomes.

| Sample | Length<br>Novoplasty | Average<br>Novoplasty<br>Coverage | Circularized | Average BWA |
| --- | --- | --- | --- | --- |
| MNHN ZO 1971-1090 | FAIL | FAIL | No | Yes (16932) |
| MNHN ZO 1971-1091 | FAIL | FAIL | No | Yes (16924) |
| MNHN ZO 1991-185 | 17164 | 1151 | Yes | No |
| MNHN ZO 1991-186 | 17296 | 2350 | No | Yes |
| MNHN ZO 1993-190 | 17174 | 878 | Yes | No |
| MNHN ZO 2017-195 | 16972 | 9640 | Yes | No |
| MNHN ZO 2017-257 | 17167 | 24765 | Yes | No |
| MNHN ZO 2017-316 | 17162-17171 | 36094 | No | Yes |
| MNHN ZO 2017-334 | 17188 | 4859 | Yes | No |
| MNHN ZO 2018-033 | 16903 | 85923 | No | Yes |
| MNHN ZO 2018-044 | FAIL | FAIL | No | Yes (16947) |
| MNHN ZO 2020-125 | 16912 | 111722 | No | Yes |

**Table S5.**
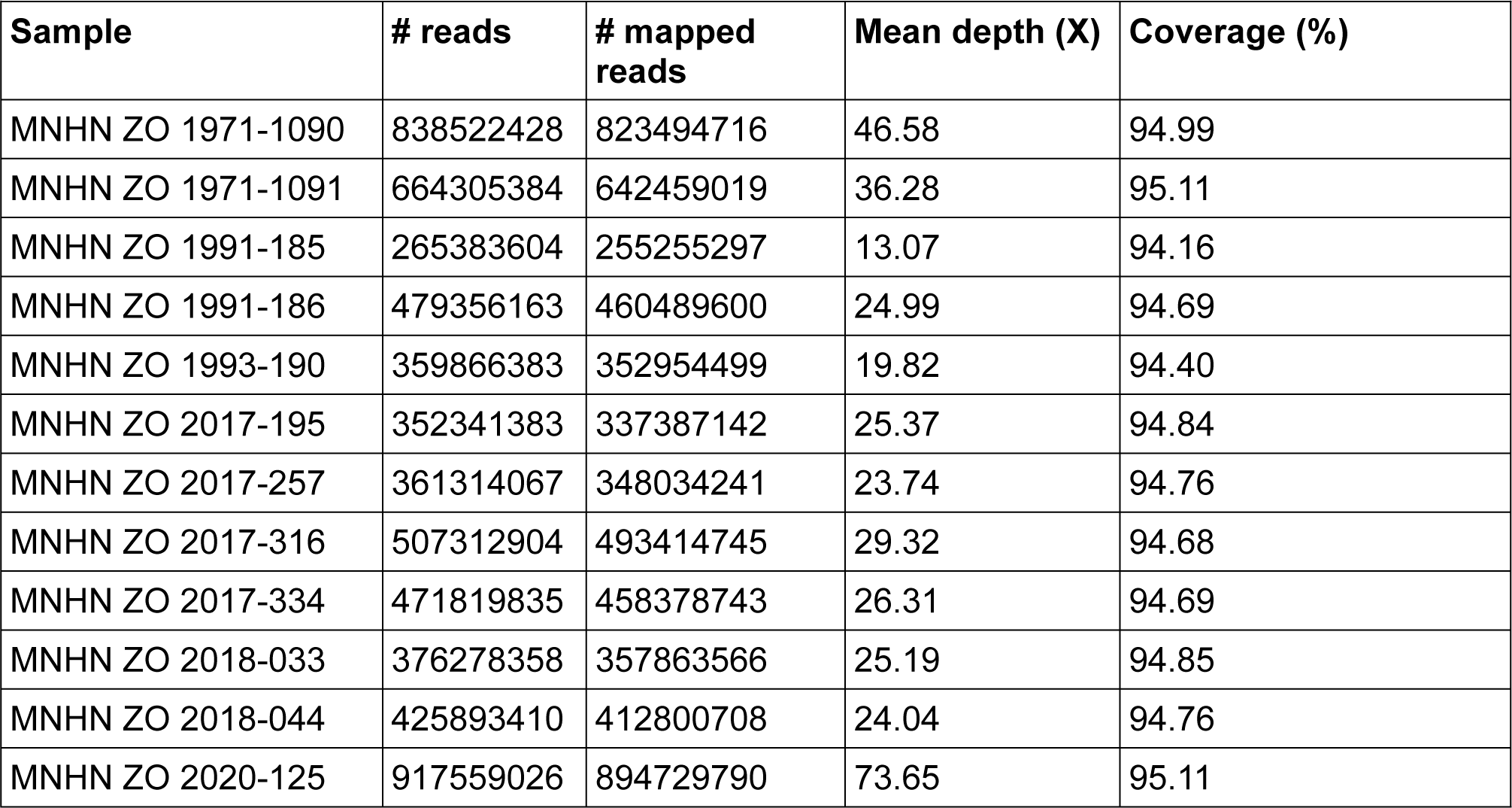
Statistics for the resequencing nuclear data, after mapping on the assembly using samtools (Danecek et al. 2021) ‘coverage’ command (v1.10)

**Table S6.** BBtools assembly statistics used to compare the assembly of *Picus viridis* in this study with that of the Northern flicker, *Colaptes auratus*.

| **Statistic** | ***Colaptes auratus*** | ***Picus viridis*** |
| --- | --- | --- |
| A | 0.2755 | 0.2727 |
| C | 0.2247 | 0.2271 |
| G | 0.2247 | 0.2272 |
| T | 0.2752 | 0.2729 |
| N | 0.0066 | 0.0000 |
| IUPAC | 0.0000 | 0.0000 |
| Other | 0.0000 | 0.0000 |
| GC | 0.4493 | 0.4544 |
| GC_stdev | 0.0778 | 0.0723 |
| Main genome scaffold total | 2369 | 47 |
| Main genome contig total | 9565 | 49 |
| Main genome scaffold sequence total | 1378.187 MB | 1228.541 MB |
| Main genome contig sequence total | 1369.051 MB<br>(0.663% gap) | 1228.526 MB<br>(0.001% gap) |
| Main genome scaffold N/L50 | 11/43.948 MB | 11/39.165 MB |
| Main genome contig N/L50 | 281/826.96 KB | 11/36.998 MB |
| Main genome scaffold N/L90 | 33/14.604 MB | 31/17.446 MB |
| Main genome contig N/L90 | 4370/50.09 KB | 32/15.416 MB |
| Max scaffold length | 117.313 MB | 107.835 MB |
| Max contig length | 15.844 MB | 107.832 MB |
| Number of scaffolds > 50 KB | 119 | 47 |
| % main genome in scaffolds > 50 KB | 97.81% | 100.00% |

**Table S7.** Number of missing callable sites for each sample after applying all the filters in the produced VCF, sorted in descending order.

| **Sample** | **Total** |
| --- | --- |
| MNHN ZO 1991-185 | 373 588 |
| MNHN ZO 2018-044 | 254 247 |
| MNHN ZO 1993-190 | 239 754 |
| MNHN ZO 1991-186 | 232 290 |
| MNHN ZO 2017-334 | 223 598 |
| MNHN ZO 2020-125 | 211 803 |
| MNHN ZO 2018-033 | 191 683 |
| MNHN ZO 2017-316 | 188 440 |
| MNHN ZO 2017-257 | 181 157 |
| MNHN ZO 1971-1091 | 175 099 |
| MNHN ZO 1971-1090 | 117 540 |
| **Total** | **2 389 199** |
| **Number of unique sites** | **848 963** |

**Table S8.** Summary of the different types of repeated elements identified using RepeatMasker.

|  | Number of elements | Length occupied | Percentage of sequence |
| --- | --- | --- | --- |
| **Retroelements** | **864793** | **316153701 bp** | **24.72 %** |
| SINEs | 1587 | 163105 bp | 0.01 % |
| Penelope | 0 | 0 bp | 0.00 % |
| LINEs | 833521 | 296643110 bp | 23.19 % |
| CRE/SLACS | 0 | 0 bp | 0.00 % |
| L2/CR1/Rex | 833521 | 296643110 bp | 23.19 % |
| R1/LOA/Jockey | 0 | 0 bp | 0.00 % |
| R2/R4/NeSL | 0 | 0 bp | 0.00 % |
| RTE/Bov-B | 0 | 0 bp | 0.00 % |
| L1/CIN4 | 0 | 0 bp | 0.00 % |
| LTR elements | 29685 | 19347486 bp | 1.51 % |
| BEL/Pao | 0 | 0 bp | 0.00 % |
| Ty1/Copia | 0 | 0 bp | 0.00 % |
| Gypsy/DIRS1 | 310 | 77693 bp | 0.01 % |
| Retroviral | 29375 | 19269793 bp | 1.51 % |
| DNA transposons | 4953 | 500460 bp | 0.04 % |
| hobo-Activator | 0 | 0 bp | 0.00 % |
| Tc1-IS630-Pogo | 0 | 0 bp | 0.00 % |
| En-Spm | 0 | 0 bp | 0.00 % |
| MuDR-IS905 | 0 | 0 bp | 0.00 % |
| PiggyBac | 0 | 0 bp | 0.00 % |
| Tourist/Harbinger | 827 | 101481 bp | 0.01 % |
| Other (Mirage, P-element, Transib) | 0 | 0 bp | 0.00 % |
| Rolling-circles | 0 | 0 bp | 0.00 % |
| Unclassified | 74215 | 36016205 bp | 2.82 % |
| **Total interspersed repeats** |  | **352670366 bp** | **27.57 %** |
| Small RNA: | 0 | 0 bp | 0.00 % |
| **Satellites** | **902** | **358374 bp** | **0.03 %** |
| **Simple repeats** | **322260** | **27250762 bp** | **2.13 %** |
| **Low complexity** | **56105** | **5021407 bp** | **0.39 %** |

**Table S9.** Summary of the different types of annotations identified using AGAT v1.2.0 (Dainat et. al 2023).

|  |  |
| --- | --- |
| **mRNA** |  |
| Number of gene | 15848 |
| Number of mrna | 16212 |
| Number of mrnas with utr both sides | 5 |
| Number of mrnas with at least one utr | 127 |
| Number of cds | 16212 |
| Number of exon | 77423 |
| Number of five_prime_utr | 73 |
| Number of intron | 61274 |
| Number of start_codon | 16119 |
| Number of stop_codon | 16136 |
| Number of three_prime_utr | 59 |
| Number of exon in cds | 77423 |
| Number of exon in five_prime_utr | 73 |
| Number of exon in three_prime_utr | 59 |
| Number of intron in cds | 61211 |
| Number of intron in exon | 61211 |
| Number of intron in intron | 48395 |
| Number gene overlapping | 73 |
| Number of single exon gene | 3315 |
| Number of single exon mrna | 3330 |
| mean mrnas per gene | 1.0 |
| mean cdss per mrna | 1.0 |
| mean exons per mrna | 4.8 |
| mean five_prime_utrs per mrna | 0.0 |
| mean introns per mrna | 3.8 |
| mean start_codons per mrna | 1.0 |
| mean stop_codons per mrna | 1.0 |
| mean three_prime_utrs per mrna | 0.0 |
| mean exons per cds | 4.8 |
| mean exons per five_prime_utr | 1.0 |
| mean exons per three_prime_utr | 1.0 |
| mean introns in cdss per mrna | 3.8 |
| mean introns in exons per mrna | 3.8 |
| mean introns in introns per mrna | 3.0 |
| Total gene length (bp) | 81357234 |
| Total mrna length (bp) | 85778659 |
| Total cds length (bp) | 16509014 |
| Total exon length (bp) | 16797198 |
| Total five_prime_utr length (bp) | 183082 |
| Total intron length (bp) | 69207345 |
| Total start_codon length (bp) | 48357 |
| Total stop_codon length (bp) | 48408 |
| Total three_prime_utr length (bp) | 105102 |
| Total intron length per cds (bp) | 68981461 |
| Total intron length per exon (bp) | 68981461 |
| Total intron length per intron (bp) | 8834961 |
| mean gene length (bp) | 5133 |
| mean mrna length (bp) | 5291 |
| mean cds length (bp) | 1018 |
| mean exon length (bp) | 216 |
| mean five_prime_utr length (bp) | 2507 |
| mean intron length (bp) | 1129 |
| mean start_codon length (bp) | 3 |
| mean stop_codon length (bp) | 3 |
| mean three_prime_utr length (bp) | 1781 |
| mean cds piece length (bp) | 213 |
| mean five_prime_utr piece length (bp) | 2507 |
| mean three_prime_utr piece length (bp) | 1781 |
| mean intron in cds length (bp) | 1126 |
| mean intron in exon length (bp) | 1126 |
| mean intron in intron length (bp) | 182 |
| Longest gene (bp) | 133036 |
| Longest mrna (bp) | 133036 |
| Longest cds (bp) | 27324 |
| Longest exon (bp) | 14959 |
| Longest five_prime_utr (bp) | 9920 |
| Longest intron (bp) | 31065 |
| Longest start_codon (bp) | 3 |
| Longest stop_codon (bp) | 3 |
| Longest three_prime_utr (bp) | 6787 |
| Longest cds piece (bp) | 14959 |
| Longest five_prime_utr piece (bp) | 9920 |
| Longest three_prime_utr piece (bp) | 6787 |
| Longest intron into cds part (bp) | 31065 |
| Longest intron into exon part (bp) | 31065 |
| Longest intron into intron part (bp) | 14959 |
| Shortest gene (bp) | 45 |
| Shortest mrna (bp) | 45 |
| Shortest cds piece (bp) | 3 |
| Shortest five_prime_utr piece (bp) | 31 |
| Shortest three_prime_utr piece (bp) | 5 |
| Shortest intron into cds part (bp) | 28 |
| Shortest intron into exon part (bp) | 28 |
| Shortest intron into intron part (bp) | 7 |
| **Without isoforms (& shortest isoforms excluded)** |  |
| Number of gene | 15848 |
| Number of mrna | 15848 |
| Number of mRNAs with utr both sides | 5 |
| Number of mRNAs with at least one utr | 126 |
| Number of cds | 15848 |
| Number of exon | 72714 |
| Number of five_prime_utr | 73 |
| Number of intron | 56928 |
| Number of start_codon | 15755 |
| Number of stop_codon | 15773 |
| Number of three_prime_utr | 58 |
| Number of exon in cds | 72714 |
| Number of exon in five_prime_utr | 73 |
| Number of exon in three_prime_utr | 58 |
| Number of intron in cds | 56866 |
| Number of intron in exon | 56866 |
| Number of intron in intron | 44398 |
| Number gene overlapping | 72 |
| Number of single exon gene | 3315 |
| Number of single exon mrna | 3315 |
| mean mrnas per gene | 1.0 |
| mean cdss per mrna | 1.0 |
| mean exons per mrna | 4.6 |
| mean five_prime_utrs per mrna | 0.0 |
| mean introns per mrna | 3.6 |
| mean start_codons per mrna | 1.0 |
| mean stop_codons per mrna | 1.0 |
| mean three_prime_utrs per mrna | 0.0 |
| mean exons per cds | 4.6 |
| mean exons per five_prime utr | 1.0 |
| mean exons per three_prime utr | 1.0 |
| mean introns in cdss per mrna | 3.6 |
| mean introns in exons per mrna | 3.6 |
| mean introns in introns per mrna | 2.8 |
| Total gene length (bp) | 81357234 |
| Total mrna length (bp) | 81290235 |
| Total cds length (bp) | 15879168 |
| Total exon length (bp) | 16164772 |
| Total five_prime utr length (bp) | 183082 |
| Total intron length (bp) | 65348767 |
| Total start_codon length (bp) | 47265 |
| Total stop_codon length (bp) | 47319 |
| Total three_prime utr length (bp) | 102522 |
| Total intron length per cds (bp) | 65125463 |
| Total intron length per exon (bp) | 65125463 |
| Total intron length per intron (bp) | 8321123 |
| mean gene length (bp) | 5133 |
| mean mrna length (bp) | 5129 |
| mean cds length (bp) | 1001 |
| mean exon length (bp) | 222 |
| mean five_prime utr length (bp) | 2507 |
| mean intron length (bp) | 1147 |
| mean start_codon length (bp) | 3 |
| mean stop_codon length (bp) | 3 |
| mean three_prime utr length (bp) | 1767 |
| mean cds piece length (bp) | 218 |
| mean five_prime utr piece length (bp) | 2507 |
| mean three_prime utr piece length (bp) | 1767 |
| mean intron in cds length (bp) | 1145 |
| mean intron in exon length (bp) | 1145 |
| mean intron in intron length (bp) | 187 |
| Longest gene (bp) | 133036 |
| Longest mrna (bp) | 133036 |
| Longest cds (bp) | 27324 |
| Longest exon (bp) | 14959 |
| Longest five_prime utr (bp) | 9920 |
| Longest intron (bp) | 31065 |
| Longest start_codon (bp) | 3 |
| Longest stop_codon (bp) | 3 |
| Longest three_prime utr (bp) | 6787 |
| Longest cds piece (bp) | 14959 |
| Longest five_prime_utr piece (bp) | 9920 |
| Longest three_prime_utr piece (bp) | 6787 |
| Longest intron into cds part (bp) | 31065 |
| Longest intron into exon part (bp) | 31065 |
| Longest intron into intron part (bp) | 14959 |
| Shortest gene (bp) | 45 |
| Shortest mrna (bp) | 45 |
| Shortest cds piece (bp) | 3 |
| Shortest five_prime_utr piece (bp) | 31 |
| Shortest three_prime_utr piece (bp) | 5 |
| Shortest intron into cds part (bp) | 28 |
| Shortest intron into exon part (bp) | 28 |
| Shortest intron into intron part (bp) | 7 |
| **Transcript** |  |
| Number of gene | 15909 |
| Number of transcript | 15995 |
| Number gene overlapping | 0 |
| Number of single exon gene | 15909 |
| mean transcripts per gene | 1.0 |
| Total gene length (bp) | 87223346 |
| Total transcript length (bp) | 82809862 |
| mean gene length (bp) | 5482 |
| mean transcript length (bp) | 5177 |
| Longest gene (bp) | 5239356 |
| Longest transcript (bp) | 133036 |
| Shortest gene (bp) | 45 |
| Shortest transcript (bp) | 45 |

## References

Achaz G. 2009. Frequency spectrum neutrality tests: one for all and all for one. Genetics. 183:249–258. doi: 10.1534/genetics.109.104042.

Anmarkrud JA, Lifjeld JT. 2017a. Complete mitochondrial genomes of eleven extinct or possibly extinct bird species. Mol Ecol Resour. 17:334–341. doi: 10.1111/1755-0998.12600.

Baudrin G et al. 2023. A Reference Genome Assembly for the Spotted Flycatcher (Muscicapa striata). Genome Biology and Evolution. 15:evad140. doi: 10.1093/gbe/evad140.

Baudry L et al. 2020. instaGRAAL: chromosome-level quality scaffolding of genomes using a proximity ligation-based scaffolder. Genome Biology. 21:148. doi: 10.1186/s13059-020-02041-z.

Bi D et al. 2019. Two new mitogenomes of Picidae (Aves, Piciformes): Sequence, structure and phylogenetic analyses. International Journal of Biological Macromolecules. 133:683–692. doi: 10.1016/j.ijbiomac.2019.04.157.

Castresana J. 2000. Selection of Conserved Blocks from Multiple Alignments for Their Use in Phylogenetic Analysis. Molecular Biology and Evolution. 17:540–552. doi: 10.1093/oxfordjournals.molbev.a026334.

Chen J et al. 2022. Characterization of the complete mitochondrial genome of the Lesser Spotted Woodpecker (Dryobates minor) and its phylogenetic position. Mitochondrial DNA Part B. 7:1504–1506. doi: 10.1080/23802359.2022.2110530.

Dainat, J., D. Hereñú, K. D. Murray, E. Davis, K. Crouch et al., 2023 AGAT: Another Gff Analysis Toolkit to handle annotations in any GTF/GFF format.

Danecek, P., A. Auton, G. Abecasis, C. A. Albers, E. Banks et al., 2011 The variant call format and VCFtools. Bioinformatics 27: 2156–2158.

Danecek, P., J. K. Bonfield, J. Liddle, J. Marshall, V. Ohan et al., 2021 Twelve years of SAMtools and BCFtools. Gigascience 10: giab008.

Dierckxsens N, Mardulyn P, Smits G. 2017. NOVOPlasty: de novo assembly of organelle genomes from whole genome data. Nucleic Acids Res. 45:e18. doi: 10.1093/nar/gkw955.

Du C, Liu LI, Liu Y, Fu Z. 2020. The complete mitochondrial genome of the Eurasian wryneck Jynx torquilla (Aves: Piciformes: Picidae) and its phylogenetic inference. Zootaxa. 4810:zootaxa.4810.2.8. doi: 10.11646/zootaxa.4810.2.8.

Duan Y, Li Y, Liang D, Shao S, Luo X. 2018. Complete mitochondrial genome of yellow-rumped honeyguide Indicator xanthonotus (Piciformes: Indicatoridae). Mitochondrial DNA Part B. 3:1278–1279. doi: 10.1080/23802359.2018.1532837.

Eo SH. 2017. Complete mitochondrial genome of white-backed woodpecker Dendrocopos leucotos (Piciformes: Picidae) and its phylogenetic position. Mitochondrial DNA Part B. 2:451–452. doi: 10.1080/23802359.2017.1357454.

Faircloth BC. 2016. PHYLUCE is a software package for the analysis of conserved genomic loci. Bioinformatics. 32:786–788. doi: 10.1093/bioinformatics/btv646.

Feng S et al. 2020. Dense sampling of bird diversity increases power of comparative genomics. Nature. 587:252–257. doi: 10.1038/s41586-020-2873-9.

Fu YX. 1995. Statistical properties of segregating sites. Theor Popul Biol. 48:172–197. doi: 10.1006/tpbi.1995.1025.

Fuchs J et al. 2008. Molecular support for a rapid cladogenesis of the woodpecker clade Malarpicini, with further insights into the genus Picus (Piciformes: Picinae). Molecular Phylogenetics and Evolution. 48:34–46. doi: 10.1016/j.ympev.2008.03.036.

Fuchs J, Pons J-M, Pasquet E, Bonillo C. 2016. Complete mitochondrial genomes of the white-browed piculet (Sasia ochracea, Picidae) and pale-billed woodpecker (Campephilus guatemalensis, Picidae). Mitochondrial DNA A DNA Mapp Seq Anal. 27:3640–3641. doi: 10.3109/19401736.2015.1079834.

Gibb GC, Kardailsky O, Kimball RT, Braun EL, Penny D. 2007. Mitochondrial genomes and avian phylogeny: complex characters and resolvability without explosive radiations. Mol Biol Evol. 24:269–280. doi: 10.1093/molbev/msl158.

Goldstein S, Beka L, Graf J, Klassen JL. 2019. Evaluation of strategies for the assembly of diverse bacterial genomes using MinION long-read sequencing. BMC Genomics. 20:23. doi: 10.1186/s12864-018-5381-7.

Hammar B. 1970. The karyotypes of thirty-one birds. Hereditas. 65:29–58. doi: 10.1111/j.1601-5223.1970.tb02306.x.

Hillier LW et al. 2004. Sequence and comparative analysis of the chicken genome provide unique perspectives on vertebrate evolution. Nature. 432:695–716. doi: 10.1038/nature03154.

Hoff KJ, Lange S, Lomsadze A, Borodovsky M, Stanke M. 2016. BRAKER1: Unsupervised RNA-Seq-Based Genome Annotation with GeneMark-ET and AUGUSTUS. Bioinformatics. 32:767–769. doi: 10.1093/bioinformatics/btv661.

Hruska JP, Manthey JD. 2021. De novo assembly of a chromosome-scale reference genome for the northern flicker Colaptes auratus. G3 (Bethesda). 11:jkaa026. doi: 10.1093/g3journal/jkaa026.

Hu Y et al. 2014. OmicCircos: A Simple-to-Use R Package for the Circular Visualization of Multidimensional Omics Data. Cancer Inform. 13:13–20. doi: 10.4137/CIN.S13495.

Issa N, Muller Y, Deceuninck B, Pons J-M, Grangé J-L. 2015. Pic vert. Atlas des oiseaux de France métropolitaine: nidification et présence hivernale. Delachaux et Niestlé.

Jarvis ED et al. 2015. Phylogenomic analyses data of the avian phylogenomics project. Gigascience. 4:4. doi: 10.1186/s13742-014-0038-1.

Katoh K, Misawa K, Kuma K, Miyata T. 2002. MAFFT: a novel method for rapid multiple sequence alignment based on fast Fourier transform. Nucleic Acids Res. 30:3059–3066. doi: 10.1093/nar/gkf436.

Kirschel ANG et al. 2021. Taxonomic revision of the Red-fronted Tinkerbird Pogoniulus pusillus (Dumont, 1816) based on molecular and phenotypic analyses. bbrc. 141:428–442. doi: 10.25226/bboc.v141i4.2021.a6.

Kolmogorov M, Yuan J, Lin Y, Pevzner PA. 2019. Assembly of long, error-prone reads using repeat graphs. Nat Biotechnol. 37:540–546. doi: 10.1038/s41587-019-0072-8.

Kozlov AM, Darriba D, Flouri T, Morel B, Stamatakis A. 2019. RAxML-NG: a fast, scalable and user-friendly tool for maximum likelihood phylogenetic inference. Bioinformatics. 35:4453–4455. doi: 10.1093/bioinformatics/btz305.

Kumar S, Stecher G, Li M, Knyaz C, Tamura K. 2018. MEGA X: Molecular Evolutionary Genetics Analysis across Computing Platforms. Molecular Biology and Evolution. 35:1547–1549. doi: 10.1093/molbev/msy096.

Kurtz S, Phillippy A, Delcher Arthur L., et al. 2004. Versatile and open software for comparing large genomes. Genome Biology. 5:R12. doi: 10.1186/gb-2004-5-2-r12.

Lai W-N, Yan S-Q, Jiao S-Y, Yao J-Y, Li Y-M. 2019. Complete mitochondrial genome of Dendrocopos canicapillus (Piciformes: Picidae). Mitochondrial DNA Part B. 4:141–142. doi: 10.1080/23802359.2018.1544051.

Librado P, Rozas J. 2009. DnaSP v5: a software for comprehensive analysis of DNA polymorphism data. Bioinformatics. 25:1451–1452. doi: 10.1093/bioinformatics/btp187.

Lomsadze A, Burns PD, Borodovsky M. 2014. Integration of mapped RNA-Seq reads into automatic training of eukaryotic gene finding algorithm. Nucleic Acids Res. 42:e119. doi: 10.1093/nar/gku557.

Lomsadze A, Ter-Hovhannisyan V, Chernoff YO, Borodovsky M. 2005. Gene identification in novel eukaryotic genomes by self-training algorithm. Nucleic Acids Res. 33:6494–6506. doi: 10.1093/nar/gki937.

Manthey JD, Moyle RG, Boissinot S. 2018. Multiple and Independent Phases of Transposable Element Amplification in the Genomes of Piciformes (Woodpeckers and Allies). Genome Biol Evol. 10:1445–1456. doi: 10.1093/gbe/evy105.

Mapleson D, Garcia Accinelli G, Kettleborough G, Wright J, Clavijo BJ. 2017. KAT: a K-mer analysis toolkit to quality control NGS datasets and genome assemblies. Bioinformatics. 33:574–576. doi: 10.1093/bioinformatics/btw663.

Mirchandani CD et al. 2023. A fast, reproducible, high-throughput variant calling workflow for evolutionary, ecological, and conservation genomics. Genomics doi: 10.1101/2023.06.22.546168.

Moreau P et al. 2018. Tridimensional infiltration of DNA viruses into the host genome shows preferential contact with active chromatin. Nat Commun. 9:4268. doi: 10.1038/s41467-018-06739-4.

Museum national d’Histoire naturelle, Office français de la biodiversité. Picus viridis Linnaeus, 1758 - Pic vert, Pivert. Inventaire National du Patrimoine Naturel. https://inpn.mnhn.fr/espece/cd_nom/3603 (Accessed July 5, 2023).

Nei M, Li WH. 1979. Mathematical model for studying genetic variation in terms of restriction endonucleases. Proceedings of the National Academy of Sciences. 76:5269–5273. doi: 10.1073/pnas.76.10.5269.

Okonechnikov K, Golosova O, Fursov M, UGENE team. 2012. Unipro UGENE: a unified bioinformatics toolkit. Bioinformatics. 28:1166–1167. doi: 10.1093/bioinformatics/bts091.

Chen Y-C, Liu T, Yu C-H, Chiang T-Y, Hwang C-C. 2013. Effects of GC Bias in Next-Generation-Sequencing Data on De Novo Genome Assembly. PLoS One. 8:e62856. doi: 10.1371/journal.pone.0062856.

de Oliveira TD et al. 2017. Genomic Organization of Repetitive DNA in Woodpeckers (Aves, Piciformes): Implications for Karyotype and ZW Sex Chromosome Differentiation. PLoS One. 12:e0169987. doi: 10.1371/journal.pone.0169987.

Paradis E. 2010. pegas: an R package for population genetics with an integrated-modular approach. Bioinformatics. 26:419–420. doi: 10.1093/bioinformatics/btp696.

Park CE et al. 2019. The complete mitochondrial genome sequence of Dendrocopos major (Aves, Piciformes, Picidae). Mitochondrial DNA Part B. 4:777–778. doi: 10.1080/23802359.2019.1565980.

Perktas U, Barrowclough GF, Groth JG. 2011. Phylogeography and species limits in the green woodpecker complex (Aves: Picidae): multiple Pleistocene refugia and range expansion across Europe and the Near East. Biological Journal of the Linnean Society. 104:710–723. doi: 10.1111/j.1095-8312.2011.01750.x.

Pfeifer B, Wittelsbürger U, Ramos-Onsins SE, Lercher MJ. 2014. PopGenome: an efficient Swiss army knife for population genomic analyses in R. Mol Biol Evol. 31:1929–1936. doi: 10.1093/molbev/msu136.

Pons J-M, Masson C, Olioso G, Fuchs J. 2019. Gene flow and genetic admixture across a secondary contact zone between two divergent lineages of the Eurasian Green Woodpecker Picus viridis. J Ornithol. 160:935–945. doi: 10.1007/s10336-019-01675-6.

Pons J-M, Olioso G, Cruaud C, Fuchs J. 2011. Phylogeography of the Eurasian green woodpecker (Picus viridis). Journal of Biogeography. 38:311–325. doi: 10.1111/j.1365-2699.2010.02401.x.

Privé F, Luu K, Vilhjálmsson BJ, Blum MGB. 2020. Performing Highly Efficient Genome Scans for Local Adaptation with R Package pcadapt Version 4. Molecular Biology and Evolution. 37:2153–2154. doi: 10.1093/molbev/msaa053.

Prum RO et al. 2015. A comprehensive phylogeny of birds (Aves) using targeted next-generation DNA sequencing. Nature. 526:569–573. doi: 10.1038/nature15697.

Shakya SB, Fuchs J, Pons J-M, Sheldon FH. 2017. Tapping the woodpecker tree for evolutionary insight. Molecular Phylogenetics and Evolution. 116:182–191. doi: 10.1016/j.ympev.2017.09.005.

Shields GF. 1982. Comparative Avian Cytogenetics: A Review. The Condor. 84:45–58. doi: 10.2307/1367820.

Simão FA, Waterhouse RM, Ioannidis P, Kriventseva EV, Zdobnov EM. 2015. BUSCO: assessing genome assembly and annotation completeness with single-copy orthologs. Bioinformatics. 31:3210–3212. doi: 10.1093/bioinformatics/btv351.

Smit A, Hubley R, Green P. 2013. RepeatMasker Open-4.0 http://www.repeatmasker.org.RMDownload.html.

Stamatakis A. 2014. RAxML version 8: a tool for phylogenetic analysis and post-analysis of large phylogenies. Bioinformatics. 30:1312–1313. doi: 10.1093/bioinformatics/btu033.

Stanke M et al. 2006. AUGUSTUS: ab initio prediction of alternative transcripts. Nucleic Acids Res. 34:W435–439. doi: 10.1093/nar/gkl200.

Tajima F. 1983. Evolutionary relationship of DNA sequences in finite populations. Genetics. 105:437–460. doi: 10.1093/genetics/105.2.437.

Talavera G, Castresana J. 2007. Improvement of phylogenies after removing divergent and ambiguously aligned blocks from protein sequence alignments. Syst Biol. 56:564–577. doi: 10.1080/10635150701472164.

Tamashiro RA et al. 2019. What are the roles of taxon sampling and model fit in tests of cyto-nuclear discordance using avian mitogenomic data? Mol Phylogenet Evol. 130:132–142. doi: 10.1016/j.ympev.2018.10.008.

Vaser R, Sovic I, Nagarajan N, Sikic M. 2017. Fast and accurate de novo genome assembly from long uncorrected reads. Genome Res. gr.214270.116. doi: 10.1101/gr.214270.116.

Walker BJ et al. 2014. Pilon: An Integrated Tool for Comprehensive Microbial Variant Detection and Genome Assembly Improvement. PLOS ONE. 9:e112963. doi: 10.1371/journal.pone.0112963.

Wiley G, Miller MJ. 2020. A Highly Contiguous Genome for the Golden-Fronted Woodpecker (Melanerpes aurifrons) via Hybrid Oxford Nanopore and Short Read Assembly. G3 (Bethesda). 10:1829–1836. doi: 10.1534/g3.120.401059.

Winnepenninckx B, Backeljau T, De Wachter R. 1993. Extraction of high molecular weight DNA from molluscs. Trends Genet. 9:407. doi: 10.1016/0168-9525(93)90102-n.

Yao J-Y et al. 2019. The complete mitochondrial genome of Picus canus (Piciformes: Picidae). Mitochondrial DNA Part B. 4:1869–1870. doi: 10.1080/23802359.2019.1614887.

Zhang G et al. 2014. Comparative genomics reveals insights into avian genome evolution and adaptation. Science. 346:1311–1320. doi: 10.1126/science.1251385.

Zhang Z, An M, Deng Y, Zhu S. 2016. The complete mitochondrial genome of the Downy woodpecker, Picoides pubescens (Piciformes: Picidae). Mitochondrial DNA A DNA Mapp Seq Anal. 27:3479–3480. doi: 10.3109/19401736.2015.1066357.

Zhou C et al. 2017. The first complete mitogenome of Picumnus innominatus (Aves, Piciformes, Picidae) and phylogenetic inference within the Picidae. Biochemical Systematics and Ecology. 70:274–282. doi: 10.1016/j.bse.2016.12.003.

